# The pericardium forms as a distinct structure during heart formation

**DOI:** 10.1101/2024.09.18.613484

**Authors:** Hannah R. Moran, Obed O. Nyarko, Rebecca O’Rourke, Ryenne-Christine K. Ching, Frederike W. Riemslagh, Brisa Peña, Alexa Burger, Carmen C. Sucharov, Christian Mosimann

**Affiliations:** Department of Pediatrics, Section of Developmental Biology, University of Colorado School of Medicine, Anschutz Medical Campus, Aurora, CO, USA; Division of Cardiology, Department of Medicine, University of Colorado Anschutz Medical Campus, Aurora, CO, USA; Cardiovascular Institute, Division of Cardiology, University of Colorado School of Medicine, Anschutz Medical Campus, Aurora, CO, USA; Bioengineering Department, University of Colorado School of Medicine, Anschutz Medical Campus, Aurora, CO, USA

## Abstract

The heart integrates diverse cell lineages into a functional unit, including the pericardium, a mesothelial sac that supports heart movement, homeostasis, and immune responses. However, despite its critical roles, the developmental origins of the pericardium remain uncertain due to disparate models. Here, using live imaging, lineage tracking, and single-cell transcriptomics in zebrafish, we find the pericardium forms within the lateral plate mesoderm from dedicated anterior mesothelial progenitors and distinct from the classic heart field. Imaging of transgenic reporters in zebrafish documents lateral plate mesoderm cells that emerge lateral of the classic heart field and among a continuous mesothelial progenitor field. Single-cell transcriptomics and trajectories of *hand2*-expressing lateral plate mesoderm reveal distinct populations of mesothelial and cardiac precursors, including pericardial precursors that are distinct from the cardiomyocyte lineage. The mesothelial gene expression signature is conserved in mammals and carries over to postnatal development. Light sheet-based live-imaging and machine learning-supported cell tracking documents that during heart tube formation, pericardial precursors that reside at the anterior edge of the heart field migrate anteriorly and medially before fusing, enclosing the embryonic heart to form a single pericardial cavity. Pericardium formation proceeds even upon genetic disruption of heart tube formation, uncoupling the two structures. Canonical Wnt/β-catenin signaling modulates pericardial cell number, resulting in a stretched pericardial epithelium with reduced cell number upon canonical Wnt inhibition. We connect the pathological expression of secreted Wnt antagonists of the SFRP family found in pediatric dilated cardiomyopathy to increased pericardial stiffness: sFRP1 in the presence of increased catecholamines causes cardiomyocyte stiffness in neonatal rats as measured by atomic force microscopy. Altogether, our data integrate pericardium formation as an independent process into heart morphogenesis and connect disrupted pericardial tissue properties such as pericardial stiffness to pediatric cardiomyopathies.

## INTRODUCTION

The formation of a functional heart requires precise integration of various cell types during embryonic development. Understanding the diverse cell lineages and mechanisms that contribute to cardiogenesis is essential to uncover the origins of congenital heart disease and to unlock the heart’s regenerative capacity. The developing heart incorporates myocardial, endocardial, and conduction system cells while being surrounded by the pericardium as a multi-layered mesothelial tissue. Covering the heart as the first functional organ in the body, the pericardium forms among the earliest organ-associated mesothelia. The functional, mature pericardium presents as a three-layered structure with i) a fibrous outer layer that in mammals attaches to the sternum and diaphragm to anchor the cardiac cavity, ii) an inner serous membrane layer that reduces friction of the constantly beating heart^1^, and iii) the epicardium as the inner-most layer that emerges from the initial pericardium to surround the myocardial surface^2–4^. The pericardium in its entirety contributes to mitigating chronic and acute stress on the heart, maintaining cardiac pressure, facilitating immune response and tissue repair, and securing and lubricating the heart within the thorax^1,5–8^. These pericardial functions become critical upon idiopathic or viral damage to the heart, including the innermost epicardium that can support signals for cardiomyocyte rearrangement and potential regeneration^9–12^. Pericardial stiffness or loss of pericardial elasticity resulting from constrictive peritoneal pressure (effusion), pericarditis, or other anomalies can impact heart function^13,14^, with the potential to also influence heart development and pediatric congenital heart anomalies^15–17^.

Mesothelia such as the pericardium have been implicated in contributing to a variety of downstream cell fates including select tissue fibroblast lineages^18–23^, yet the lineage connectivity and clonality of mesothelial progenitor cells remain vastly unknown. How pericardial precursors resolve amongst cardiac progenitors and the developed heart, as well as how the unique tissue properties of the pericardial layers compared to the heart are set up during development, also awaits resolution. Cardiac progenitors emerge as bilateral field within the lateral plate mesoderm (LPM), the dedicated mesodermal progenitor domain that forms at the lateral edge of the post-gastrulation embryo^24–26^. The vertebrate endocardium and cardiomyocytes emerge from at least two distinguishable LPM territories known as the first heart field (FHF) and second heart field (SHF)^27–34^. In mammals, the FHF primarily forms the left ventricle myocardium to support systemic circulation, while the SHF forms the atria and the right ventricle driving lung circulation^25^. In mice, a juxta-cardiac field (JCF) has been implicated as a transcriptionally distinct population contributing to parts of the ventricle and epicardium^35^. Additionally, the heart forms within a broader Tbx1-expressing progenitor domain termed the cardiopharyngeal field (CPF) that is chiefly defined to include progenitors for the myocardium and distinct branchiomeric muscles in head and neck^36–44^. However, the molecular, developmental, evolutionary, and conceptual relationships of these heart-contributing populations, as well as their comparison across species, remain to be fully explored. Among the heart-contributing lineages, the origins of the pericardium have received notably little attention, in part due to missing developmental concepts of how mesothelial lineages relate to, and develop with, their associated organs.

Within the anterior LPM (ALPM), FHF cells migrate medially to form the linear heart tube that extends with later-differentiating SHF progenitors^25,45^. The pericardium has been challenging to study due to its dynamic development and lack of genetic markers^46^. Lineage tracing studies in mouse and zebrafish firmly established that the pericardium emerges among the ALPM^19^, with cell labeling-based observations proposing a SHF or JCF origin of the pericardium^35,47–50^. Myocardial SHF progenitors contributing to the outflow tract closely associate with the dorsal pericardial wall in mammals, forming among a continuum with pericardial progenitors before rupture of the dorsal mesocardial connection^51,52^. However, the position, lineage relationship, and migration trajectory of initial pericardial progenitors relative to the heart remain to be defined. Early mesothelial origins have been defined using zebrafish and mouse as lateral-most, Hand2-expressing LPM^19^. Single-cell transcriptomics in zebrafish, mouse, and embryonic stem cells have revealed several LPM progenitor lineages that feature high expression of Hand2 or of its paralog Hand1 and co-expression of distinct transcription factor combinations^19,35,50^, underlining the heterogeneity and complexity among Hand1/2-positive LPM progenitors. Genetic lineage tracing and live imaging of *hand2:EGFP* transgenic zebrafish revealed the complex migration of visceral versus parietal mesothelial layers including of the pericardium. Notably, pericardium/epicardium-contributing progenitors including the JCF have been described to express *Hand* genes^19,35,50,53^. Mechanistically, pioneering work in Xenopus has linked canonical Wnt/β-catenin signaling to balancing pericardial versus myocardial differentiation, with active signaling supporting proliferation of the former while suppressing the latter^54–56^. Modulating canonical Wnt/β-catenin signaling is applied in the *in vitro*-driven derivation of pericardial and later epicardial lineages with potential utility in therapeutic applications^57–60^. Collectively, and in addition to its mesothelial anatomy, these results suggest that the pericardium follows a unique morphogenetic trajectory among heart-contributing lineages.

Here, by combining live-imaging, machine learning-based cell tracking, single cell transcriptome-based trajectories, and functional studies in zebrafish, we resolve the earliest origins of the pericardium. We contextualize pericardial precursors to reside among the continuous, *hand2*-positive mesothelium-forming LPM progenitors. Emerging lateral to, and outside of, the well-described heart tube-forming cardiac field, our data define pericardial progenitors as distinct mesothelial and not cardiac lineage *per se* with a unique transcriptional signature that is conserved in mammals. We show that canonical Wnt signaling is required to establish proper pericardial morphogenesis and tissue tension. Lastly, we link serum exposure to the Wnt antagonist sFRP1 as found in pediatric dilated cardiomyopathy (DCM) to reduced pericardial elasticity in neonatal rats, revealing a possibly contributing factor to disease progression. Our findings expand our concepts of how the pericardium integrates with the heart to form a single functional organ system, providing a paradigm for how mesothelia integrate with, and affect, their associated organs during development and disease.

## RESULTS

### The pericardium forms continuous with the mesothelium-forming LPM

The BAC-based transgenic *hand2:EGFP* zebrafish line recapitulates endogenous *hand2* expression in LPM-derived cell types including the myocardium, endocardium, pectoral fins, mesothelium, and pharyngeal arches^19,61^. While imaging mesothelial progenitors *in vivo*, we previously documented pericardial progenitors among *hand2:EGFP* transgene-expressing cells in developing zebrafish embryos by their progressive lateral, downward migration over the anterior yolk^19^ (**Fig. 1A,B**). Notably, *hand2*-expressing parietal mesothelial progenitors migrate laterally outward, while other LPM lineages such as the heart or endothelial progenitors migrate towards the midline^19^. To gain spatio-temporal insights into how and exactly where from the pericardium emerges during cardiac development, we revisited *hand2:EGFP* expression by imaging together with several transgenic zebrafish reporters that delineate cardiac lineages *in vivo*.

**Fig. 1:**
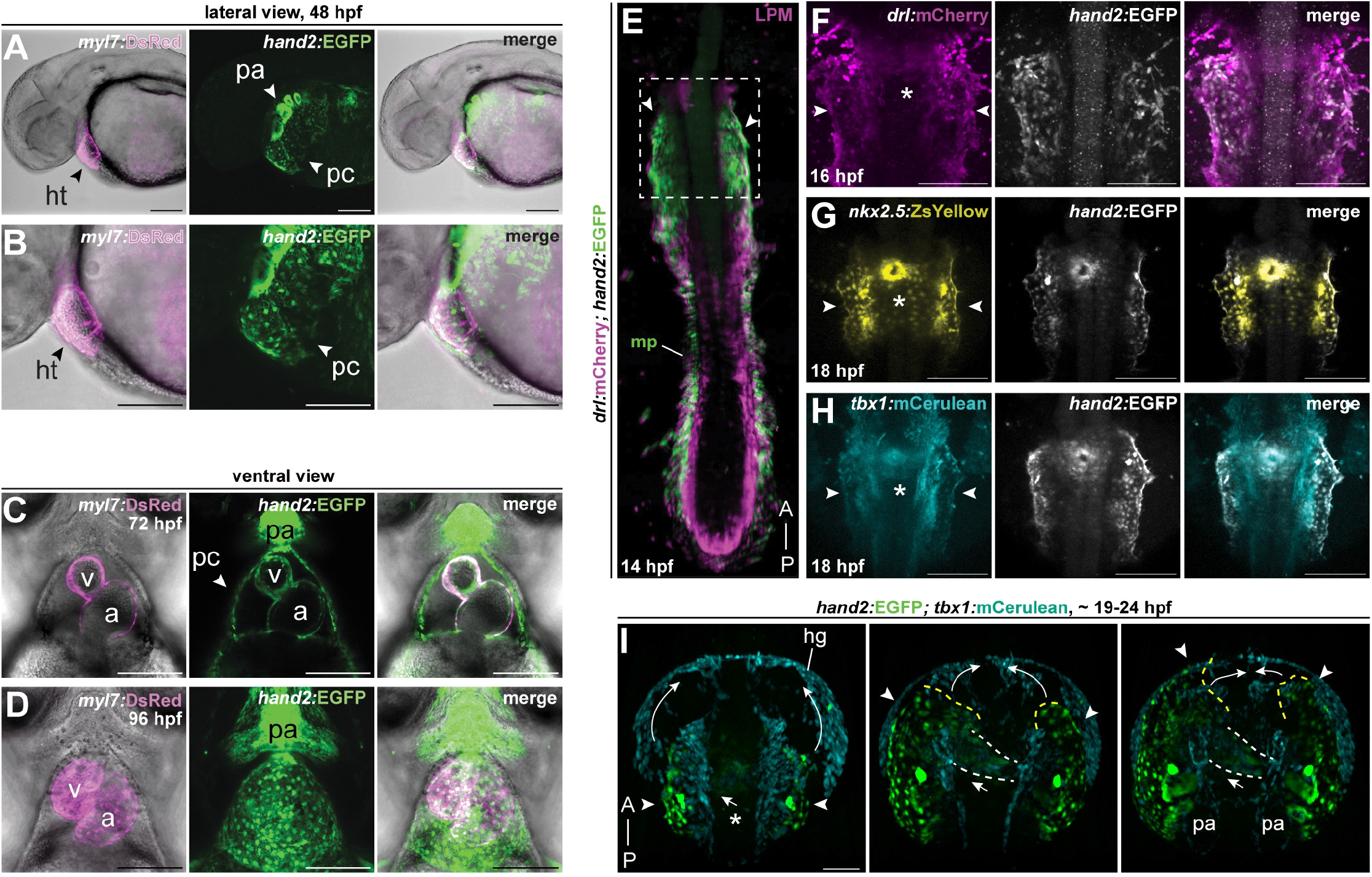
The pericardium forms continuous with the mesothelium-forming LPM. **A-D**. Anatomy of the heart and pericardium in zebrafish embryos and early larvae. **A,B**. Lateral confocal imaging of transgenic *hand2:EGFP;myl7:DsRed* zebrafish, anterior to the left. Embryo showing *hand2:EGFP* and *myl7:DsRed* (myocardium marker) co-expression in the heart tube and *hand2:EGFP-*expressing cell populations in the pericardium, posterior mesothelium, pectoral fin, and pharyngeal arches at 48 hpf (**A**, 10x; **B**, 20x). **C,D**. Ventral confocal imaging (**C** 72 hpf single Z-slice, **D** 96 hpf max projection, anterior to the top) of *hand2:EGFP;myl7:DsRed* embryo showing reporter co-expression in the atrium and ventricle myocardium and *hand2:EGFP-*expressing cells in the pharyngeal arches and pericardial sac surrounding the heart (**C**), with the pericardium acquiring a mesh-like squamous epithelial structure at 96 hpf (**D**). **E**. SPIM-based Mercator projection of double-transgenic *hand2:EGFP;drl:mCherry* embryo at 14 hpf, anterior to the top, showing the anterior-to-posterior extent of lateral-most *hand2:EGFP*-expressing LPM cells fated as mesothelial progenitors (mp), with the anterior-most extent of the bilateral stripes indicated by white arrowheads. Dotted box outlines region of interest for subsequent imaging panels. **F-H**. Dorsal confocal imaging of transgenic reporter combinations at the level of the heart field, anterior to the top, asterisks showing the midline where the heart field converges, arrowheads at lateral mesh-like cells. **F**. *hand2:EGFP;drl:mCherry* embryo at 16 hpf prior to medial heart field migration, showing *hand2:EGFP*-expressing cell populations comprising lateral-most LPM. **G**) *hand2:EGFP;nkx2*.*5:ZsYellow* embryo with dual-marked emerging cardiac disk. **H**. *hand2:EGFP;tbx1:mCerulean* embryo with similar co-expression across bilateral mesh-like cells and the cardiac disk. **I**. SPIM-based live imaging stills from Movie 1 depicting a *hand2:EGFP;tbx1:mCerulean* embryo from 19-24 hpf, anterior to the top, showing *hand2:EGFP-*expressing lateral-most cells migrating away from the midline towards the front (migration front, yellow dashed line), while cardiac precursors form the heart tube towards the midline (white dashed line). White arrows for migration direction. pa, pharyngeal arches; pc, pericardium; ht, heart; v, ventricle; a, atrium; hg, hatching gland. Scale bars **A-D** 100 μm, **E-H** 200 μm, **I** 50 μm.

As an anatomical overview, we first crossed *hand2:EGFP* to *myl7:DsRed*^63^ that exclusively labels cardiomyocytes upon differentiation past 18 hours post-fertilization (hpf)^92^ (**Fig. 1 A-D**). While *myl7* and *hand2* reporters co-expressed in the medially converging and looping myocardium of both ventricle and atrium (**Fig. 1 C-D**), more lateral *hand2:EGFP*-positive;*myl7:dsRed*-negative cells migrated laterally over the yolk as an epithelial sheet that enclosed the developing heart tube^19^ (**Fig. 1 A-B**). These observations indicate that *hand2:EGFP*-positive ALPM cells migrate laterally to the medially converging heart field to form the pericardium.

In the dynamic heart field, expression of the pan-LPM marker *drl:mCherry* gradually confines from broader LPM activity to restricted expression in FHF-derived lineages and in the endocardium^62,93^. We previously established that the lateral-most *hand2*;*drl* double-positive cell population fated for mesothelia forms a continuous band of cells along the circumference of the developing embryo^19^: *hand2:EGFP*-expressing cells outline the lateral edge of the LPM over the entire anterior-to-posterior axis of the embryo, including the ALPM lateral to the emerging heart field (**Fig. 1E**). In double-transgenic *hand2:EGFP;drl:mCherry* embryos at 16-18 hpf, we confirmed co-expression of *hand2* and *drl* reporters in the cardiac progenitors condensing towards the midline and in the developing cardiac disc, as well as once again in the more lateral cells, confirming these as LPM-derived (**Fig. 1F**). This architecture, together with their subsequent lateral migration^19^ (**Fig 1A-B**), matches the localization of prospective mesothelial progenitors within the LPM.

*Nkx2*.*5* is an evolutionarily conserved homeobox-containing transcription factor gene involved in cardiomyocyte differentiation and is expressed in cardiac progenitors of both the first and second heart fields^94–96^. Consistent with prior descriptions^97,98^, we observed co-expression of *nkx2*.*5:ZsYellow*^33^ and *hand2:EGFP* in cells localized to the bilateral territories of myocardial progenitors at 18 hpf, prior to their migration to the midline and subsequently in the developing cardiac disc (**Fig. 1G**). This pattern is consistent with the joint contribution of Nkx2.5 and Hand2 in establishing the linear heart tube and in later cardiomyocyte formation^65,99,100^. In addition to myocardial progenitors, we also observed expression of *nkx2*.*5:ZsYellow* in the lateral-most ALPM progenitors that are co-labeled by *hand2:EGFP* as a part of the continuous mesothelial progenitor stripe (**Fig. 1E,G**).

*tbx1*-based reporters label the deeply conserved cardiopharyngeal field (CPF) among the ALPM that form the myocardial and branchiomeric muscle groups^69,101–103^. In addition to the expected dual *tbx1:mCerulean*;*hand2;EGFP* reporter labeling of heart tube progenitors, we again observed *tbx1:mCerulean* and *hand2:EGFP* reporter expression colocalizing in cells within the lateral-most, anterior bilateral domains next to the developing heart tube (**Fig. 1H**). Taken together, our imaging comparison indicates that several markers of the ALPM and specifically cardiac progenitors are expressed in both the prospective bilateral heart field that forms the heart tube, as well as the more laterally located cells that form a mesh-like epithelial architecture past 18 hpf along the embryo margin.

From our reporter expression analysis and considering early LPM architecture^26^, we hypothesized that the presumptive pericardial cells are the anterior-most extension of the continuous mesothelial progenitors surrounding the developing zebrafish embryo^19^, and positioned more lateral to the emerging myocardial and endocardial progenitors (**Fig. 1E**). This model is consistent with, and contextually expands, mouse-based findings of a juxta-cardiac position attributed to epicardium-contributing (and thus pericardium-derived) LPM progenitors^35^ and Hand1/2 expression in pericardial progenitors^50^. We corroborated our hypothesis by live imaging *tbx1:mCerulean*;*hand2:EGFP* embryos using selective plane illumination microscopy (SPIM or light sheet microscopy)^19,69,104^. Time lapse imaging of dorsal views revealed the anterior-most *hand2:EGFP*-expressing cells that are continuous with the mesothelial cell band migrating as bilateral fields anteriorly and then merging to form the pericardium (**Fig. 1I**). Concurrently, the heart tube progenitors migrate medially before jogging to the left; the elongating primitive heart tube extends into the field surrounded by the enveloping migration of the pericardial progenitors (**Fig. 1I, Supplementary Movie 1**). Notably, the pericardial *hand2:EGFP* progenitors migrate asymmetrically, with the left progenitors migrating further beyond the midline before fusing with the less-advancing right side (**Fig. 1I, Supplementary Movie 1**).

Taken together, our multi-reporter imaging in zebrafish reveals that ALPM-derived pericardial progenitors share the expression of key markers assigned to heart tube progenitors, and that pericardial progenitors form as the anterior-most extension of a continuous *hand2:EGFP*-labeled mesothelial LPM progenitor field at the lateral edge of the early embryo.

### Pericardial progenitors have distinct migratory trajectories

To corroborate our hypothesis on the distinct origin and dynamics of pericardial progenitors in relation to the emerging heart tube, we performed light sheet-based live imaging of developing *hand2:EGFP* zebrafish embryos as pericardial progenitors migrate over the yolk (**Fig. 1I, Fig. 2A-F**). By applying a multi-sample imaging and processing workflow^86^, we captured the development of heart tube and pericardial progenitors over the course of cardiogenesis from 20 to 60 hpf. Defining the final fates and positions of pericardial versus myocardial cells at the imaging end point, we sought to back-track^87^ the individual lineages to establish their original arrangement within the ALPM. Additionally, to extract quantitative morphometric data, we applied the machine learning-based Object Classification and Lineage tools in Imaris (see Methods for details) (**Fig. 2B-F, Supplementary Movies 2,3, Supplementary Fig. 1A**). Around 20 hpf of zebrafish development, bilateral cardiac progenitors coalesce as cardiac disk and begin heart tube extrusion^34,45,105^, while mesothelial precursors have initiated their lateral migration over the yolk^19^ (**Fig. 2B, Supplementary Movies 2,3**). To chart the cellular dynamics of each population, we continued to follow this process throughout cardiogenesis until the pericardium has well-encapsulated the functional, beating heart at 60 hpf (**Fig. 2B-F**).

**Fig. 2:**
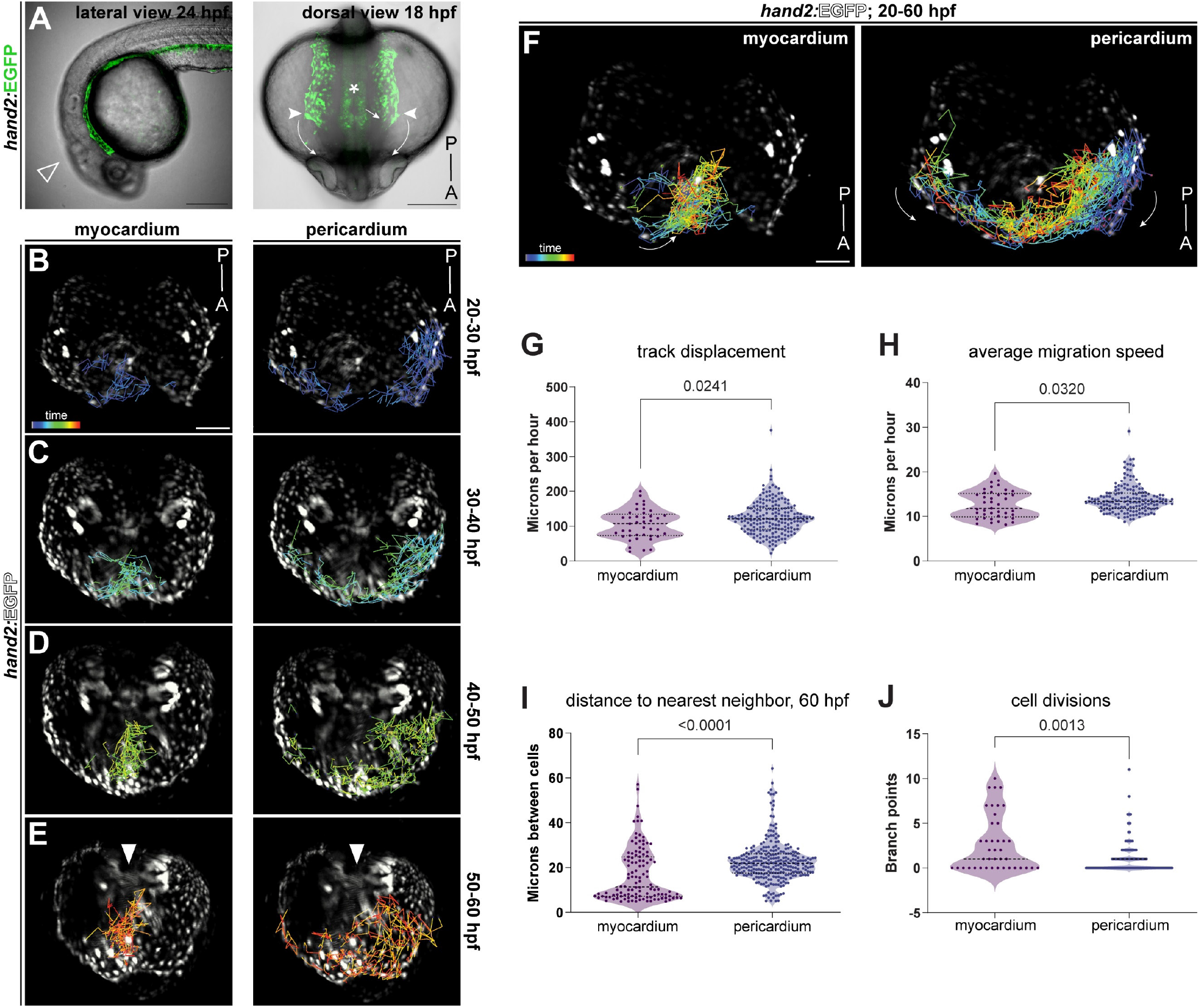
Pericardial progenitors have distinct migratory trajectories among heart-contributing lineages. **A**. Representative lateral (24 hpf, anterior to the left) and dorsal (18 hpf, anterior to the bottom) view of *hand2:EGFP* zebrafish embryos depicting embryo orientation and direction of pericardial migration as oriented in subsequent stills. **B-E** Still frames of SPIM-based timelapse movie showing *hand2:EGFP* cell trajectories as determined by machine learning-based backtracking, anterior to the bottom. The representative stills cover development from 20-30 hpf (**B**), 30-40 hpf (**C**), 40-50 hpf (**D**), and 50-60 hpf (**E**). Arrowheads in **E** show jitter from heartbeat. **F**. Snapshot of timelapse movie (Movies 2,3, anterior to the bottom) showing summarized *hand2:EGFP* cell trajectories of the developing myocardium (left) and pericardium (right) from 20-60 hpf. **G-J**. Quantification of tracked cells across replicates for myocardial and pericardial trajectories. **G**. Track displacement from position start over the time series in the myocardium and pericardium. **H**. Quantification of the speed of individual cell tracks over time. **I**. Quantification of distance between cells (nearest neighbor) at 60 hpf. **J**. Quantification of number of branch points, where a single track branches into a new track indicating a cell division, in the myocardium and pericardium. **G-J** analyzed using Mann-Whitney *t-*test, N=2 independent *in toto* tracking series. Scale bar **A** 100 μm **B-F** 70 μm.

In our resulting timelapse imaging, we once more observed pericardial precursors emerging as a mesh-like territory at the outer lateral-most edge of the heart field (**Fig. 2B-F**), as described above (**Fig. 1E-I**). Notably, early pericardial progenitors migrate with trajectories and dynamics distinct from medial heart field migration and heart tube elongation: while the cardiac progenitors migrate to the midline inward from the bilateral ALPM, the pericardium forms by asymmetric, lateral, and anterior migration of the mesh-like lateral epithelium (**Fig. 2F, Supplementary Movies 2,3**).

Imaris-based lineage mapping and machine learning off our imaging datasets further quantified the differences in cellular properties of both the pericardium and heart tube (see Methods for details). Pericardial cells travel a further distance from their original position as they migrate as bilateral coherent units over the yolk of the embryo, whereas cardiac precursors maintain their position near the midline throughout cardiogenesis (**Fig. 2G**). Compared to cardiac progenitors, pericardial progenitor cells also travel at a greater average speed throughout cardiogenesis as they encapsulate the developing heart tube (**Fig. 2H**). By 60 hpf, when both pericardium and heart are well-functional, pericardial precursors have passed through fewer cell divisions than cardiac precursors and show a greater distance between neighboring cells compared to the densely packed heart tube (**Fig. 2J,I**). Together, our timelapse imaging and cell tracking documents distinct migratory trajectories and cellular behaviors between the forming heart tube and the pericardium, consistent with the latter forming as mesothelial entity.

### Pericardium progenitors emerge among mesothelial lineages

We next sought to probe whether pericardial progenitors are a transcriptionally distinct cell population in the emerging LPM and what gene expression signature defines them. We conducted a 10xGenomics-based single-cell RNA-sequencing (scRNA-seq) analysis of zebrafish LPM cells at 10 hpf (tailbud stage, end of gastrulation) when the LPM has coalesced into its bilateral stripe architecture (**Fig. 3A**). We dissociated *drl:mCherry*;*hand2:EGFP* zebrafish embryos and isolated all mCherry-expressing cells to capture the broader, *drl*-marked LPM that includes the *hand2:EGFP*-expressing lateral plate lineages (**SFig. 2A**). We sequenced the transcriptomes of 3,305 *drl:mCherry*-labeled cells that constitute about 6.6% of all cells in the zebrafish embryo at tailbud stage, in line with previous work (**SFig. 2B**)^19,93^. Transcripts for *EGFP* in the *hand2*-positive LPM provided additional anchors for cell identities and specificity. We performed clustering analysis using the Seurat 5 R-Package^75^ to segregate captured cells into 18 principal clusters based on previously established gene expression patterns, along with marker genes identified through differential expression analysis (**Fig. 3B, SFig. 3**) (see Methods for details)^19,75,93^.

**Fig. 3:**
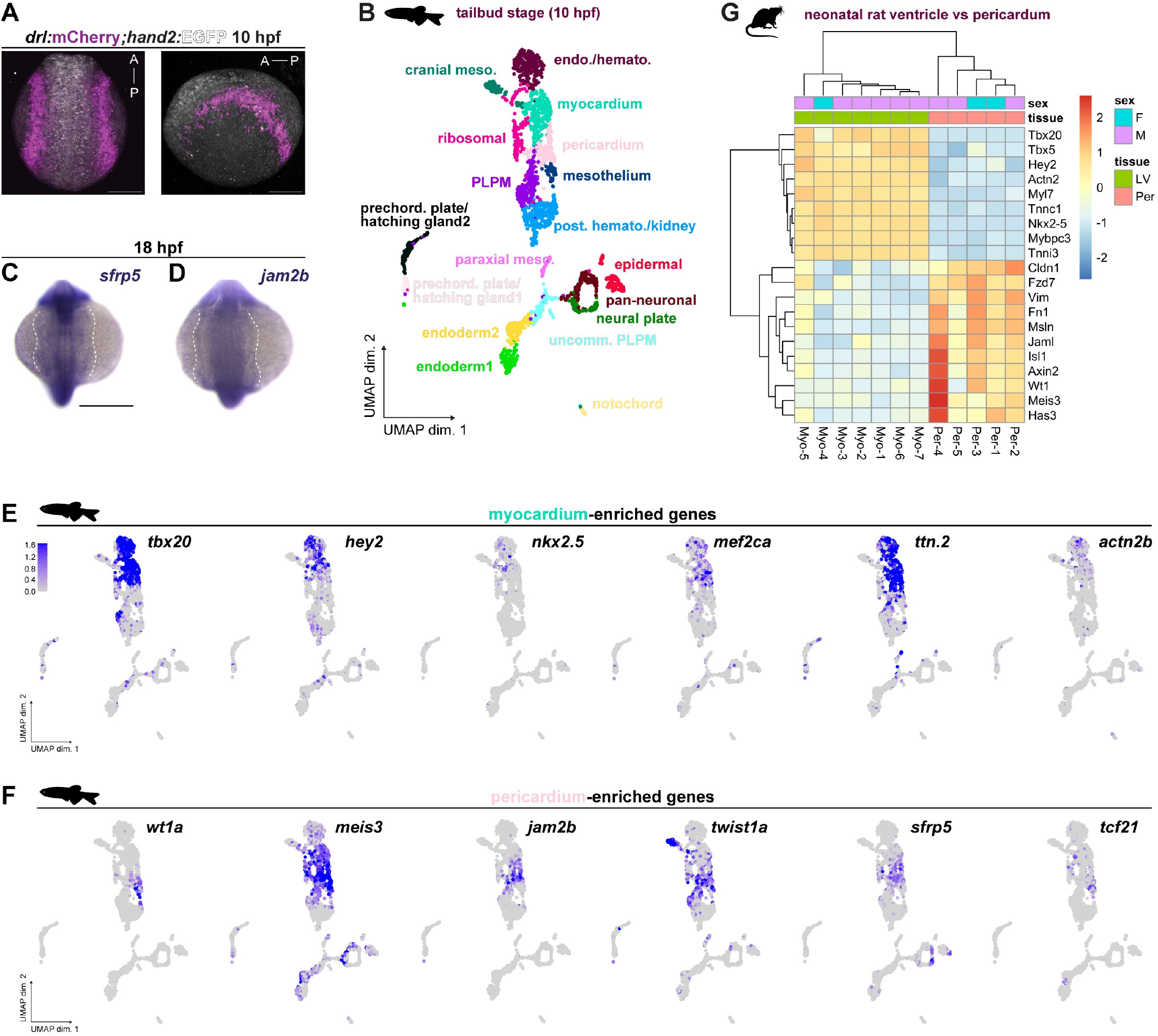
Pericardial and myocardial precursors are transcriptionally distinct populations. **A**. Representative confocal max projection of *drl:mCherry*;*hand2:EGFP* double-transgenic embryo at 10 hpf as used for FACS-based isolation of post-gastrulation LPM for 10xGenomics-based single cell transcriptomics; anterior-posterior axis as indicated. **B**. UMAP plot of single cell transcriptomes of mCherry-sorted 10 hpf *drl:mCherry*;*hand2:EGFP* zebrafish embryo cells showing 18 significant cell clusters, colored by identified subpopulation. **C,D**. Whole-mount mRNA *in situ* hybridization of representative pericardial/mesothelial genes *sfrp5* (**C**) and *jam2b* (**D**), white dashed outlines to emphasize mesh-like spread of the expression domains lateral to the embryo axis; anterior to the top. **E,F**. UMAP plots of key myocardial (**E**) and pericardial genes (**F**) expressed across identified cluster identities. Cell representations are colored by scaled expression values using lower and upper 2%-quantiles as boundaries. **G**. Clustering analysis of bulk mRNA-sequenced left ventricle myocardium (Myo) and pericardium (Per) from neonatal rats for genes defining myocardium versus pericardium as identified from the zebrafish-based scRNA-seq analysis in **B**. Heatmap bins colored by row-scaled log2-normalized counts; columns (samples) split by tissue type; rows and columns ordered by hierarchical clustering (scaled expression values), sex of sample indicated on top. Scale bar **A,B** 200 μm.

The resulting annotated clusters predominantly represent LPM-derived cell types including broadly cranial mesoderm, hemangioblasts and kidney precursors, myeloid and hematopoietic precursors, cardiomyocytes, mesothelia, as well as hatching gland cells and undifferentiated mesoderm (**Fig. 3B**). One cluster was positive for designated cranial mesoderm progenitor markers^19,41,93,106,107^, namely *fsta, foxc1a, tbx1*, and *meox1*. Hemangioblasts and kidney precursors were characterized by the expression of *lbx2, fli1a*, and *pax2a*, in line with our previous data showing intermingled lineage origins of these LPM derivatives^19,108^. Myeloid and hematopoietic precursors were positive for specified markers *lmo2, npas4l, tal1*, and *gfi1ab*^*108–111*^. Myocardial precursors were defined by the expression of *tbx20, mef2ca, hey2, actn2b, ttn*.*2*, and *nkx2*.*5*^*65*,*97*,*112*,*113*^. In contrast, broadly defined mesothelial progenitors expressed genes including *tcf21, sema3aa, wt1a, meis3, adra1d*, and *kank1a* as distinct gene expression signature^19,108,114^. Additionally, we uncovered enrichment of endodermal markers *sox17, foxa2, foxa3* in two select clusters^19,106,107^ (**Fig. 3B**). The presence of endoderm aligns with previous findings that cells positive for the *drl* reporter at tailbud stage constitute a combination of separate endoderm- and LPM-primed populations^19,81^.

Among the resolved *hand2*-positive mesothelial cells, we identified a transcriptionally distinct cluster of *hand2*-positive cells that through markers with previously described mRNA *in situ* hybridization patterns correspond to lateral-most cells of the early bilateral ALPM; this region coincides with our observed *hand2:EGFP*-positive cells migrating over the yolk to form the pericardium^19^ (**Fig. 3C-D**). In addition to *nkx2*.*5* as captured with our reporter imaging (**Fig. 1G**), mRNA *in situs* for several genes enriched in this ALPM cluster show expression patterns laterally and beyond the heart field, forming a dispersed, mesh-like pattern over the yolk during somitogenesis; specifically, *jam2b* and *sfrp5*^19,79,115,116^ (**Fig. 3C-D**). *sfrp5* in particular delineates the outermost cells of the ALPM and the remaining mesothelium (**Fig. 3C**). This mesh-like pattern is evident in the endogenous expression of *hand2* and transgenic reporters including *drl:mCherry, hand2:EGFP*, and others active within the ALPM (**Fig. 1E-H**). Combined with our live imaging data, we postulate that this select ALPM cluster represents pericardial progenitors.

We next asked if this differential transcriptome signature between the myocardium and pericardium is also detectable in mammalian systems and remains in more mature hearts. We compiled cluster-defining LPM gene lists for myocardium versus mesothelium/pericardium as informed from our zebrafish scRNA-seq (**Fig. 3B**) and prior work^19,108^. We then compared the expression of these select gene lists to bulk RNA-sequencing of isolated pericardia (n=5) and left ventricle myocardia (n=7) from neonatal 3-week-old rats (*Rattus norvegicus*) (**Fig. 3G-F**). Overall, the neonatal pericardium differentially expressed 1712 genes (11.5%) compared to the sequenced ventricular myocardium (**Fig. 3G**). Comparing the myocardium to mesothelium/pericardium-defining genes, the sequenced neonatal rat samples hierarchically separated into distinct myocardial and mesothelial/pericardial expression signatures (**Fig. 3G**). These signatures were independent of sex differences and included genes across different functional classes associated with tissue properties specific to each tissue type (**Fig. 3G**). Examples include cardiac transcription factor and myofibril genes in the myocardium, while the pericardium enriched for collagens, extracellular matrix, and signaling components related to the Wnt signaling cascade (**Fig. 3G-F**). Together, despite sharing the expression of several genes associated with ALPM and cardiopharyngeal progenitors, our comparative work reveals a distinct gene signature consistent with pericardial emergence from the continuous mesothelium.

### Pericardial progenitors follow a distinct differentiation trajectory

Our transcriptome analysis resulted in gene sets assigned to myocardial, pericardial, and more broadly mesothelial progenitors at the end of gastrulation. Pericardial marker genes include *jam2b, sfrp5, tmem88b, nr2f1a, meis2b* and *twist1a*, which are also enriched in the mesothelium-primed LPM^19^ (**Fig. 4A**). In addition, this cluster also retained expression signatures typical of cardiomyocyte precursors, including *gata5, gata6, nkx2*.*5*, and *ttn*.*2* ^*65,97,112,113*^ (**Fig. 4A**). Notably, we once more found shared, albeit heterogeneous, expression of classic cardiac genes in both the pericardial and myocardial cluster, in line with imaging data and their shared emergence in the ALPM (**Fig. 1E-H, Fig. 4B**). *nkx2*.*5* reporter activity exemplifies this heterogeneity^33,98^, with stronger reporter activity in the more medial putative myocardial progenitors compared to the spread-out, more lateral pericardial progenitors (**Fig. 4B**).

**Fig. 4:**
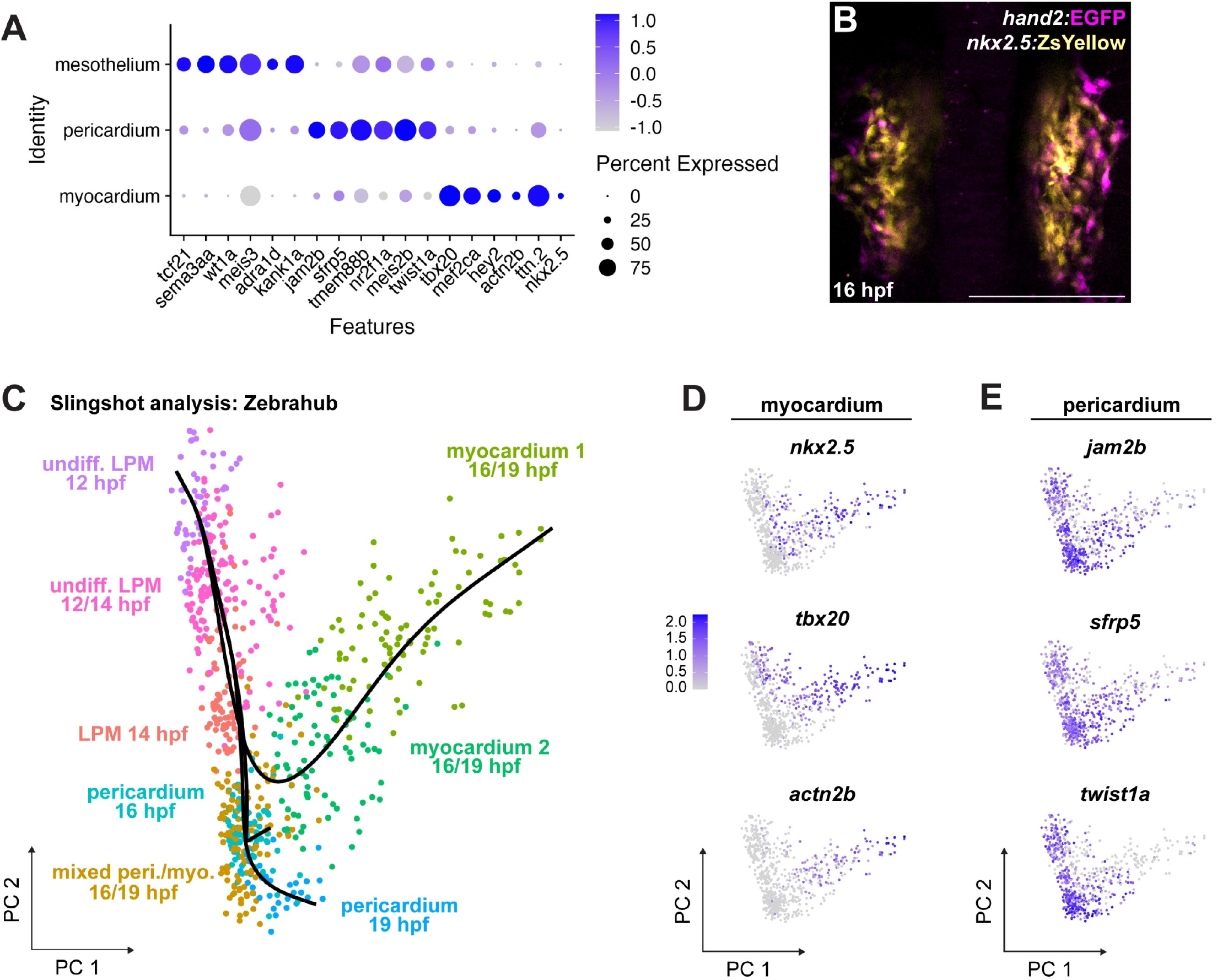
The pericardial lineage trajectory becomes distinct prior to heart tube formation. **A**. Dotplot including key cell fate marker genes to annotate broad mesothelial, pericardial, and myocardial clusters, respectively. Dots colored by column-scaled mean expression (log-transformed library-size-normalized counts) and sized by expression frequency (fraction of cells with non-zero counts). **B**. Dorsal confocal imaging of representative dual-transgenic *hand2:EGFP;nkx2*.*5:ZsYellow* zebrafish embryo at 16 hpf showing marker co-expression and heterogeneity in prospective cardiac and pericardial progenitor cells around the heart field; anterior to the top. **C-E** Slingshot-based trajectory inference analysis of early LPM cells assigned using key marker genes to the public Zebrahub data set of single cell transcriptomes throughout zebrafish development. Inferred end points for myocardium and pericardium indicated as color-coded clusters (**C**). PCA plots of key myocardial (**D**) and pericardial genes (**E**) expressed across identities over time. Cells are colored by scaled expression values using lower and upper 2%-quantiles as boundaries. Scale bar **B** 200 μm.

To independently corroborate pericardial origins from mesothelial instead of classic heart field progenitors, we sought to apply our cluster-defining gene lists for single cell-based trajectory inference. This approach requires a comprehensive scRNA-seq dataset with sufficient cells and multiple timepoints to compute pseudotemporal reconstruction of lineage differentiation^117–120^. To perform developmental trajectory inference analysis using Slingshot^77^, we transposed our LPM cluster definitions for cardiomyocytes and putative pericardial progenitors onto the publicly available multi-timepoint, whole-embryo scRNA-seq data from the Zebrahub repository^76^. The complete Zebrahub data covers 10 hpf to 10 dpf zebrafish sequencing data^76^, including our developmental time points of interest.

Our curated gene list matched with the Zebrahub cells designated as “primitive heart tube cells” or heart” from 12 hpf, 14 hpf, 16 hpf, and 19 hpf, which we then selected for re-analysis as UMAP containing pericardium, myocardium, and early precursor clusters (see Methods for details) (**SFig. 4A-D**). This re-analysis once more documented myocardial and putative pericardial lineages as distinct cell clusters upon their progressive specification in the LPM (**Fig. 4C, SFig. 4E-G**). We then reduced the principal component analysis of this subset of Zebrahub cells to generate lineage trajectories using Slingshot (**Fig. 4C, Fig. S4**). This trajectory inference from a starting point of more naïve, undifferentiated LPM in late gastrulation throughout somitogenesis plotted a rapidly diverging differentiation trajectory between the myocardium and pericardium (**Fig. 4C-E**), following expression dynamics of the key pericardial and myocardial-defining genes we had defined (**Fig. 4A, SFig. 5A-B**). Together, this orthogonal analysis further supports a model that within the ALPM, pericardial progenitors diverge from heart-primed early on and behave akin to mesothelial progenitors.

### The pericardium forms independently of the heart tube

The concerted development of mesothelia with their associated organs, and how their formation is influenced by organ-specific anomalies, remain understudied^22,23,121,122^. We therefore sought to determine how genetic perturbations affecting heart development influence pericardium formation.

The perturbation of *mef2ca* and *mef2cb* allows cardiomyocytes to be specified but arrests their differentiation, and double-morphant zebrafish fail to form a functional heart^71,123^. At 24 hpf, in uninjected *hand2:EGFP* control embryos, the heart tube has formed at the midline, with pericardial precursors retaining their mesh-like, lateral position prior to their anterior migration to envelop the embryonic heart (**Fig. 5A**). At 60 hpf, cardiac looping has occurred and both atrium and ventricle are clearly discernible by *hand2:EGFP* and anti-MHC staining (MF20), while *hand2:EGFP*-positive pericardial cells border the primitive heart tube (**Fig. 5B**). In contrast, *mef2ca/b* double-morphant *hand2:EGFP* zebrafish failed to form a heart tube by 24 hpf, while pericardial progenitors remained present and comparable to wildtype in morphology and position (**Fig. 5C**). Consistent with previous findings^71,123^, *mef2ca/b* double-morphants did not form hearts with organized myocardial structure and fail to beat at 36 hpf (**Fig. 5D**). Nonetheless, the emerging pericardial sac retained its characteristic structure that encapsulates the under-developed heart tube (**Fig. 5D**). These observations indicate that pericardium emergence is not dependent on the timing or fidelity of heart tube formation, consistent with the pericardium forming as a mesothelial tissue during cardiogenesis.

**Fig. 5:**
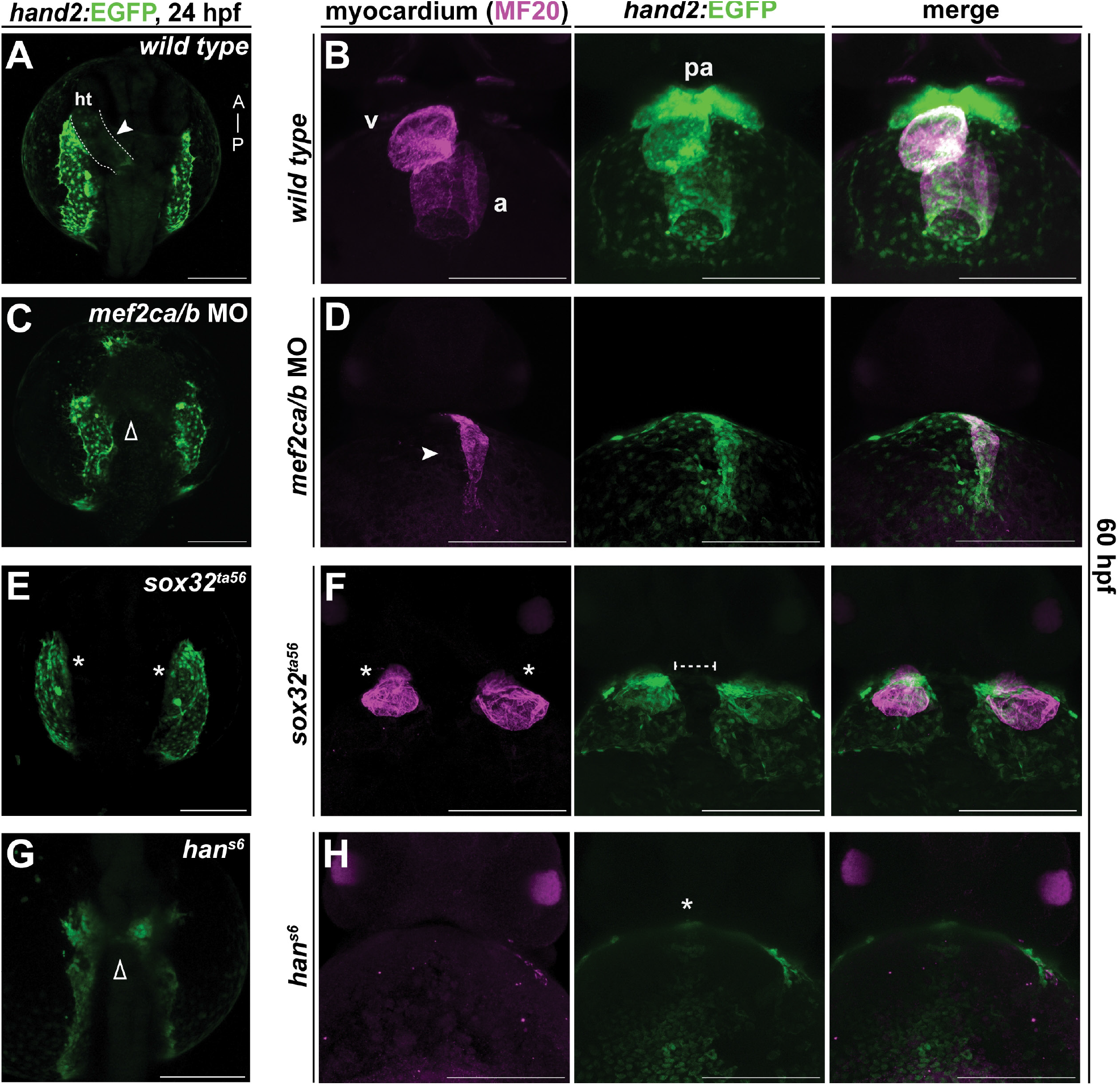
The pericardium forms despite developmental insults to heart formation. **A-H**. Confocal imaging of *hand2:EGFP* transgenic zebrafish (green), live (**A,C,E,G**, 24 hpf dorsal views, anterior to top) or with immunofluorescence using anti-MHC antibody (MF20, magenta) to show myocardium (**B,D,F,H**, 60 hpf ventral views, anterior to top). **A**. Representative confocal image of 24 hpf *hand2:EGFP* transgenic embryo showing *hand2:EGFP*-expressing wild-type pericardial precursors and heart tube development (white arrowhead, dashed outline) as dorsal view. **B**. Wild-type reference for *hand2:EGFP*-expressing embryos and MHC counterstained at 60 hpf. **C,D**. Delayed and disrupted heart tube formation upon mef2ca/b knockdown still allows for pericardium formation. Representative confocal image of 24 hpf *hand2:EGFP* embryo injected with both *mef2ca* and *mef2cb* morpholinos (**C**), showing *hand2:EGFP*-expressing pericardial precursors and severely delayed or absent medial migration of heart tube progenitors (open arrowhead) and at 60 hpf with rudimentary heart tube (**D**, white arrowhead). **E,F**. Loss of endoderm and cardia bifida still allows for pericardium formation. Representative confocal image of 24 hpf *sox32*^*ta56*^-homozygous mutant zebrafish in the *hand2:EGFP* background with pericardial precursors present and absent midline migration of heart tube progenitors (**E**); cardia bifida (*casanova* phenotype) and two pericardial cavities formed at 60 hpf (**F**). **G,H**. Representative confocal image of 24 hpf *han*^*s6*^ mutant zebrafish in the *hand2:EGFP* background showing disrupted pericardial and cardiac precursor migration (open arrowhead) (**G**) and absent cardiac chambers (asterisk) and a pericardial cavity at 60 hpf (**H**). v, ventricle; a, atrium. Scale bar **A-H** 200 μm.

The midline migration of heart tube progenitors requires the adjacent endoderm^124–128^. Zebrafish mutants and morphants for the key endoderm transcription factor gene *sox32* fail to form all endoderm and develop cardia bifida (*casanova, cas*), the formation of two hearts due to failed midline convergence^66,127,129,130^. In *sox32*^*ta56*^ (*cas*) mutant embryos displaying classic cardia bifida (**Fig. 5E,F**), we invariantly found that each heart was encapsulated by a separate pericardium of variable size (**Fig. 5F**). The two pericardia formed an interface at the midline and did not mix, forming two separate pericardial cavities by 60 hpf (**Fig. 5F**). We conclude that while endoderm integrity is required for proper heart and pericardium migration, the pericardium still forms upon perturbed midline convergence of the heart tube progenitors. This observation proposes that an endoderm-derived signal might be required to merge the left and right pericardial primordia that migrate beyond the anterior extent of the endoderm (**Fig. 1H**).

We next assessed the previously established zebrafish *hand2* mutant *han*^*S6*^ (*hands off*) that harbors a presumptive *hand2* null allele^65^. *han*^*S6*^-mutant zebrafish exhibit blistering and shrinking of the ventral fin fold along with an uneven distribution of mesothelial progenitors^19,82^, in addition to the well-documented cardiac, pharyngeal, and pectoral fin defects^65,131^. As previously established^65,132^, at 24 hpf, the bilateral ALPM of *han*^*S6*^ zebrafish is much narrower when compared to wildtype siblings, with few observable pericardial precursors spreading out as an epithelial mesh (**Fig. 5G**). At 60 hpf, *hand2* mutants lack MF20-positive cardiomyocytes, and presented with few detectable pericardial cells without a formed pericardium (**Fig. 5H**), consistent with *hand2* mutants featuring perturbed mesothelial progenitor migration and reduced cell number^19^. Together, our data documents that genetic perturbations that interrupt cardiac progenitor migration (*sox32*) or heart tube formation and differentiation (*mef2c*), but that do not cause broader mesothelial defects (*hand2*), still allow for formation of a heart-surrounding pericardium.

### Wnt/β-catenin signaling is active in pericardial progenitors

Our transcriptomics analysis provides an opportunity to probe molecular mechanisms driving the unique migration and tissue properties of the forming pericardium. We applied Metascape analysis^73^ to identify statistically enriched functional annotations of the top-50 differentially expressed genes in the pericardium cluster and cardiomyocyte cluster from our scRNA-seq, respectively (**Fig. 3,4**). Metascape output for cardiomyocyte precursors enriched for anticipated gene ontology terms associated with differentiating, specializing cells including tube morphogenesis and heart morphogenesis (**Fig. 6A**). Metascape analysis of the top-50 genes from the pericardium cluster in contrast highlighted migrating epithelia-associated processes, in line with pericardial epithelial migration and morphogenesis (**Fig. 6B**). In addition, the pericardium output included the gene ontology term “regulation of canonical Wnt signaling pathway” as the only developmental signaling pathway with significant enrichment (**Fig. 6B**). The canonical Wnt signaling pathway controls cell fate, migration, and proliferation in diverse developmental contexts, including throughout cardiogenesis through gene regulation by the nuclear β-catenin-TCF/LEF complex^133–135^. In *Xenopus*, canonical Wnt signaling has been linked to initially support the proliferation of pericardial precursors and to inhibit myocardial proliferation, requiring tight regulation to restrict the Wnt ligand range and signaling activity between the heart and pericardium^54,136–138^. Despite these insights, the tissue-level influence of canonical Wnt signaling on pericardium formation remains unclear.

**Fig. 6:**
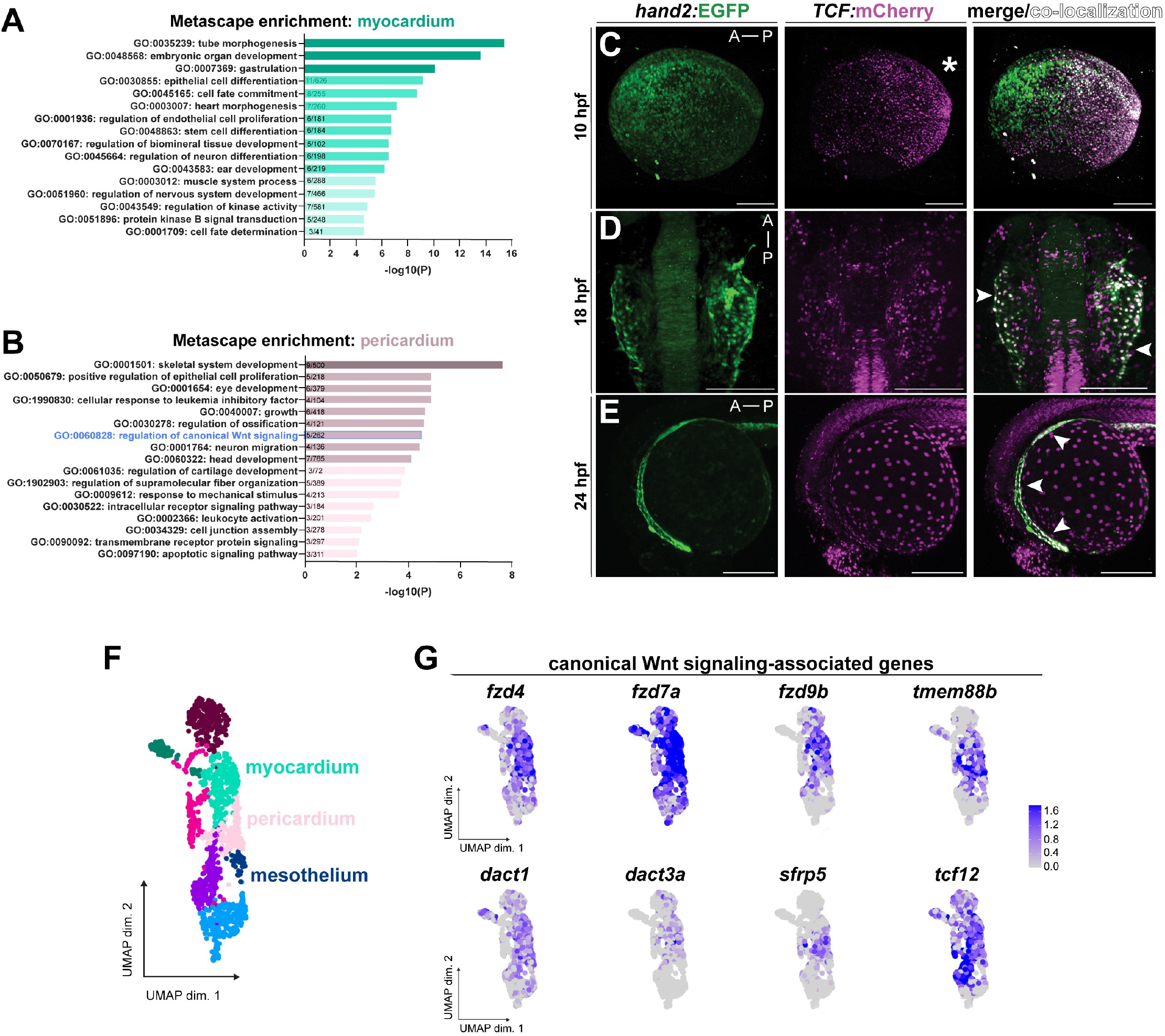
Wnt/β-catenin signaling is differentially active across pericardial progenitors. **A,B**. Metascape analysis of top-50 cluster-defining genes from early LPM scRNA-seq; gene ontology terms enriched in myocardial cluster (**A**), and pericardial cluster (**B**); note regulation of canonical Wnt signaling as significant in pericardial cells. **C-E**. Confocal max projections of representative transgene expression for *hand2:EGFP* (green) and *7xTCF:mCherry* (magenta, broadly reading out canonical Wnt signaling activity) with colocalization (white overlay), embryonic axis as indicated. At 10 hpf, canonical Wnt signaling shows a graded activity from the posterior towards the anterior (**C**, lateral view, asterisk). At the start of heart field convergence, various anterior LPM cells show heterogeneous canonical Wnt activity, with lateral-most putative pericardial progenitors strongly expressing the *7xTCF:mCherry* reporter (**D**, dorsal view, arrowheads). This pattern continues throughout to 24 hpf (**E**, lateral view, white arrowheads). **F,G**. Expression of canonical Wnt signaling-associated genes across myocardial and pericardial/mesothelial cells at tail bud stage in zebrafish. Cropped UMAP plot to depict cell clusters of interest (**F**) and individual Wnt signaling-associated genes, with expressing cells colored by scaled expression values using lower and upper 2%-quantiles as boundaries (**G**). Scale bar **C-E** 200 μm.

To visualize active canonical Wnt/β-catenin signaling in developing zebrafish embryos, we used the synthetic reporter-based *7xTCF-Xla*.*Siam:nlsmCherry*^*ia5*^ (abbreviated as *7xTCF:mCherry)* transgenic line that reads out β-catenin-TCF/LEF activity including during heart formation^64,139^. We crossed *7xTCF:mCherry* to *hand2:EGFP* to visualize any colocalization of Wnt/β-catenin activity in the *hand2:EGFP*-expressing pericardial precursors. Upon imaging at tailbud stage (10 hpf), initially faint *hand2:EGFP* expression appears in a mild intensity gradient concentrated at the anterior of the embryo in the presumptive heart field-forming ALPM (**Fig. 6C**). The canonical Wnt-stimulated *7xTCF:mCherry* reporter instead is more concentrated at the posterior of the embryo (**Fig. 6C**), in line with the classic contribution of canonical Wnt/β-catenin signaling to establishing posterior structures^64,140–143^. At 18 hpf, in the layer of the ALPM with the migrating heart field and mesh-like pericardial progenitors, we observed double-positive cells densely colocalized in the lateral-most edge of the heart field where pericardial progenitors reside, with more sparse colocalization at the midline where the presumptive heart tube is converging (**Fig. 6D**). At 24 hpf, as *hand2:EGFP*-positive pericardial cells migrate over the yolk of the embryo to form the pericardial cavity, they co-express *7xTCF:mCherry* primarily at the leading edge of the migrating epithelial sheet (**Fig. 6E**). This activity pattern continued in pericardial progenitors during their migration as they encapsulate the developing heart tube (**Fig. 6E**). These imaging data indicate that Wnt/β-catenin signaling is continuously active in pericardial progenitors throughout pericardium differentiation and migration. Wnt/β-catenin signaling-associated genes encoding pathway components and target genes also prominently featured within our tailbud stage scRNA-seq dataset across myocardial and pericardial/mesothelial cell clusters (**Fig. 6F-G**). Together, our observations align with previous findings in Xenopus that implicated high levels of canonical Wnt signaling in the pericardium to promote pericardial proliferation^54,136^.

### Canonical Wnt signaling influences pericardial cell density

Building on previous and our new data, we next aimed to test what impact canonical Wnt/β-catenin signaling has on the morphogenesis and functionality of the pericardium. To inhibit Wnt signaling during pericardium formation, we used the chemical compound IWR-1 that increases β-catenin degradation through Axin stabilization^144^. As control for influences of heart function on pericardium formation, we also treated separate embryo cohorts with 2,3-Butanedione monoxime (BDM) that inhibits myosin ATPase and stops heartbeat^145^; BDM treatment enabled us to compare pericardial phenotypes that resulted from selectively disrupting heart function, as canonical Wnt signaling is a major regulator of pacemaker formation^146,147^. We treated *hand2:EGFP;myl7:DsRed* embryos with a vehicle control (DMSO, n=11), IWR-1 (n=10), and BDM (n=9), respectively at 18 hpf. Treatment at this timepoint circumvents gross influences on embryo patterning by Wnt inhibition and aligns with the onset of medial migration of heart tube progenitors and anterior-lateral migration of pericardial progenitors^34,148,149^ (**Fig. 1E-I**). At 72 hpf, we then imaged the pericardial cavity of *hand2:EGFP;myl7:DsRed* larvae to compare size and density of cells in IWR-1- or BDM-treated pericardia, respectively, to controls treated with DMSO vehicle only (n=6 cells analyzed per sample) (**Fig. 7A-B**).

**Fig. 7:**
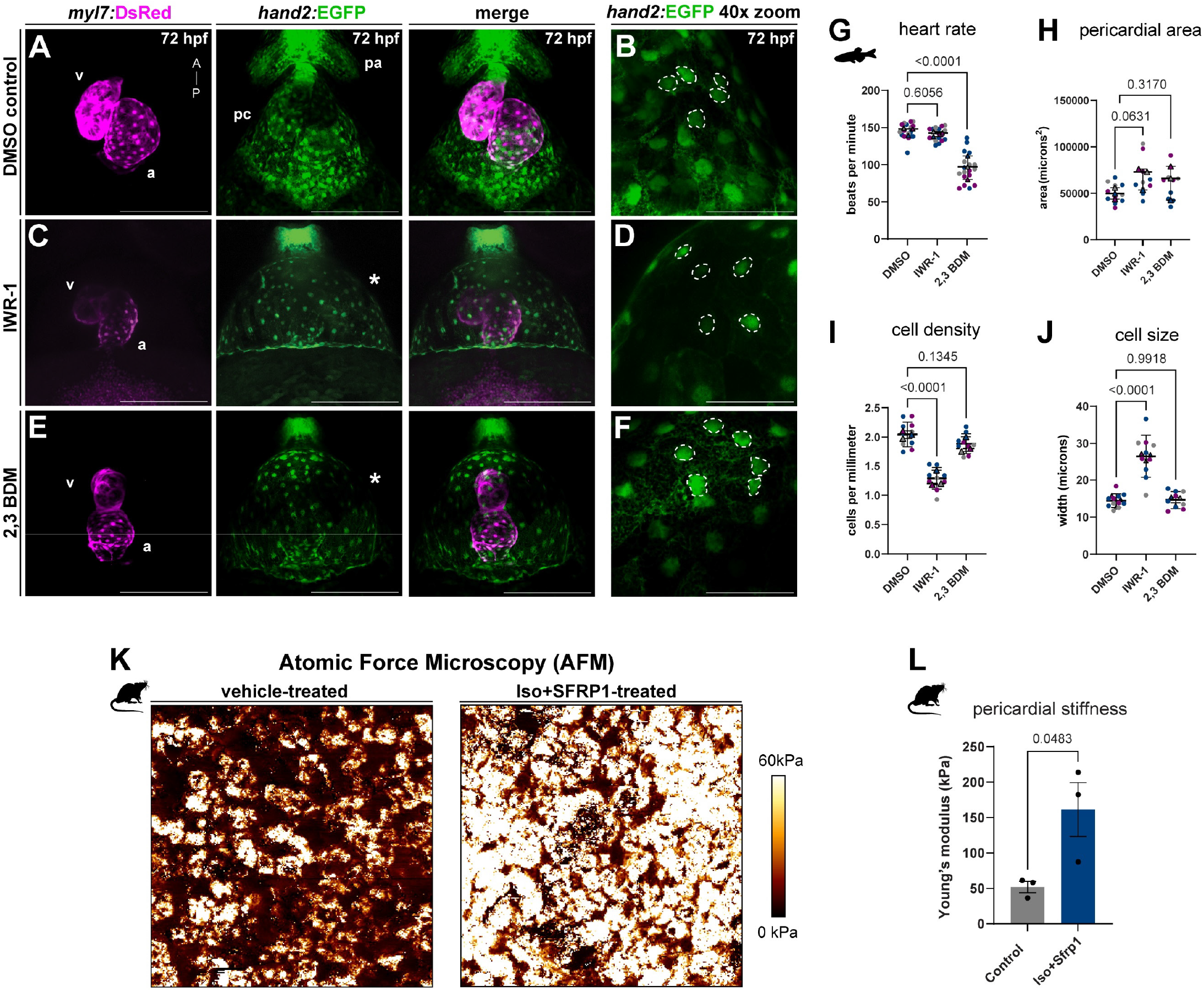
Inhibition of Wnt/β-catenin alters pericardial morphology and stiffness. **A-E**. Ventral confocal imaging (max projection) of representative 72 hpf *hand2:EGFP;myl7DsRed* larvae undergoing distinct treatments as indicated; anterior to the top; v, ventricle; a, atrium. **A,B** Representative larvae treated with DMSO vehicle only as control at 18 hpf overnight showing *hand2:EGFP-*expressing pericardial sac surrounding the larval zebrafish heart at 72 hpf (**A**, 20x) and cellular density (**B**, 40x zoom, representative nuclei marked with dashed lines). **C,D**. Ventral images of representative *hand2:EGFP;myl7:DsRed* larvae treated with the Wnt signaling inhibitor IWR-1 at 18 hpf overnight, showing expanded pericardial sac and edema with large, stretched cells surrounding the larval zebrafish heart at 72 hpf (**C**, 20x) and lower cellular density (**D**, 40x). **E,F**. Ventral images of *hand2:EGFP;myl7:DsRed* larvae treated with BDM as myosin-II inhibitor at 18 hpf overnight showing expanded pericardial sac with normal cell distribution in the larval zebrafish heart at 72 hpf (**E**, 20x) and cellular density (**F**, 40x). **G-J**. Quantifications of pericardial and cardiac features following the diverse treatments. One-way ANOVA, n=6 embryos over 3 independent experiments. **G**. Heart rate of vehicle-treated, Wnt-inhibited, and myosin II-inhibited (2,3 BDM) embryos. **H**. Pericardial area (overall distribution as per ventral view), showing increased pericardial area in IWR-1 treated larvae. **I**. Cell density (cells per square millimeter), showing decreased cell density in IWR-1-treated larvae only. **J**. Cell size showing increases in IWR-1-treated embryos only. **K,L**. Increased tissue stiffness in the pericardium of rats treated with PBS (control) or combined Isoproterenol (Iso) and SFRP1 (n=3). Neonatal rats (0-to-4-day old rats) were injected intraperitoneally with 0.05mg/kg/day of human recombinant sFRP1 protein and Iso in an animal model of pediatric dilated cardiomyopathy. Atomic force microscopy (AFM) of dissected pericardia provided measures for Young’s modulus (kPa) as readout for tissue elasticity, with treated pericardia showing increased stiffness (**K**) as quantified per sample(**L**, unpaired 2-tailed *t*-test, p = 0.0482). Each dot represents an individual sample. Representative images of a control and iso+sfrp1 treated rats. Scale bar **A,C,E** 200 μm, **B,D,F** (40x) 50 μm.

Wnt-inhibited embryos featured severe pericardial edema with minimal to no overt changes to gross morphology of the myocardium (**Fig. 7C-D**). IWR-1-treated embryos also showed limited changes in heart rate, indicating that our systemic Wnt inhibition at 18 hpf has minimal overt effects on cardiac function (**Fig. 7G**). In contrast, BDM-treated embryos showed the anticipated decreased heart rate and also pericardial edema, likely due to reduced blood flow and disrupted osmoregulation (**Fig. 7E-G**). In IWR-1-treated embryos, the pericardial cells appeared enlarged and expanded, whereas both the BDM- and DMSO-treated animals maintained a comparable cell size even when edema had developed (**Fig. 7H,J**). Quantification of the cell density in the observable pericardial area revealed reduced pericardial cell density and numbers in IWR1-treated larvae, in line with the expanded and stretched individual cells (**Fig. 7H-I**). Connecting to prior work linking canonical Wnt/β-catenin activity to pericardial proliferation^54^, we conclude that while still permissive to pericardium formation, reduced Wnt/β-catenin pathway activity results in a pericardium with fewer cells while covering a comparable area to wild type.

### SFRP1 exposure alters pericardial stiffness in neonatal rats

Our finding that reduced canonical Wnt/β-catenin signaling results in a pericardium with fewer cells led us to speculate that tissue rigidity or elasticity may be altered as a result of the changed cellular density. Pediatric patients with dilated cardiomyopathy (DCM) have been found to harbor increased levels of the Wnt-inhibiting SFRP1 protein in blood circulation^150^ with as-of-yet unclear onset of SFRP1 expression, its source, or causal influence on the DCM phenotype. Our previous work established that exposing cultured neonatal rat ventricular cardiomyocytes (NRVMs) to SFRP1 increased their cellular stiffness^150^. Extrapolating from our observations in zebrafish and the conservation of a maintained pericardial gene program in neonatal rats (**Fig. 3G**), we reasoned that chronic inhibition of canonical Wnt signaling by circulating sFRP1 might influence the rigidity of the mammalian pericardium.

As a general hallmark, catecholamines (such as Isoproterenol, Iso) are increased in serum of pediatric heart failure patients and contribute to increasing disease severity^151,152^. We therefore devised an injection-based approach to mimic aspects of pediatric dilated cardiomyopathy in neonatal rats through exposure to Iso and recombinant SFRP1. We intra-peritoneally injected 0.1 mg/kg/day of Iso and 0.05 mg/kg/day of recombinant SFRP1 protein every other day for the first 5 days, and then every day for 7 days, to recapitulate increasing catecholamine and SFRP1 levels in human DCM serum (see Methods for details). We next isolated the pericardium of Iso+SFRP1 -treated rats and of vehicle-treated controls and performed atomic force microscopy (AFM) to measure tissue stiffness by determining Young’s modulus (expressed in kPa) (**Fig. 7K**). While AFM of wild-type pericardia consistently returned a Young’s modulus of 36-61 kPa (mean 52.03 +/-13.65) (**Fig. 7K,L**), Iso+SFRP1-treated neonatal rat pericardia showed a consistent increase in tissue stiffness ranging from 87-213 kPa (mean 161.2 +/-65.85) (N=3 per condition) (**Fig. 7K,L**). Together, these findings indicate that modulation of canonical Wnt/ β-catenin signaling through persistent exposure to secreted antagonists influences pericardial cell composition and tissue rigidity in development and in pathological conditions.

## DISCUSSION

Despite the central contributions of the pericardium and of its derivative epicardium to cardiac homeostasis and disease, the developmental origins of the pericardium have remained challenging to grasp. Our results uncover that the pericardium develops as a mesothelial lineage that is transcriptionally and by lineage distinct from classic heart tube progenitors. Consolidating disparate models of pericardium origins relative to heart field progenitors, our findings advance our concept of how mesothelia co-develop together with their associated organs. Our work also extends the current understanding of cell fate complexity within the ALPM that includes the heart field and integrates LPM-derived mesothelial progenitors into cardiogenesis. Our live imaging and tracking highlight the emergence of the pericardium as the anterior-most portion of a continuous mesothelial band, expanding the developmental context of earlier, pioneering observations of heart tube formation using transgenic reporters and live labeling in zebrafish, chick, and mouse^19,34,69,153–157^. The existence of a lateral-most mesothelial progenitor band also evokes a potentially ancestral embryonic architecture in early chordates where LPM-derived mesothelial progenitors establish visceral support by migrating laterally within the coelom prior to organogenesis^81,158^.

Traditional concepts of cardiogenesis have assigned heart-contributing lineages to the first or second heart field^30,34,47,50,157,159^. Individual studies have proposed that pericardial progenitors might originate from the SHF, expanding more recently to include the so-called juxta-cardiac field and Hand1-positive progenitors, which are linked to epicardial origins in mice^35,52^. Additionally, the ALPM includes *tbx1*-marked progenitors that contribute to both myocardial and craniofacial muscle development, combined into the cardiopharyngeal field or mesoderm^36– 38,160,161^. Our data now clarifies these models by defining the pericardium as a mesothelial and not cardiac lineage *per se*. Pericardial progenitors share key ALPM marker genes with myocardial progenitors by virtue of their shared LPM origin and anterior position^9,19,35,153,154,162,163^ (**Fig. 1,3,4, SFig.3-5**). Nonetheless, their lineage origin and developmental trajectory are distinct from the heart tube and align with a continuous mesothelial band at the embryo’s edge^19^ (**Fig. 1E, Fig. 4C,D**). This model integrates heart and pericardium formation into a functional unit akin to endoderm-derived internal organs with associated LPM-derived mesothelia^21,22,164^. Notably, in mammals, the dorsal portion of the heart tube initially forms continuous with the dorsal pericardial wall and subsequently separates^51,52,165^. This contrasts the situation in zebrafish where the heart extrudes from the initial cardiac disk and elongates with concomitant pericardial progenitor migration^34,69,105^ (**Fig. 1, 2**). This seemingly disparate morphology can be reconciled by regarding the junction of myocardium and dorsal pericardial wall as the earlier-forming boundary between cardiomyocyte versus mesothelial/pericardial stripes during medial-to-lateral fate patterning in the post-gastrulation ALPM^19,52^. Challenging to observe *in vivo* due to tissue depth and folding, the remnant SHF cells embedded in the dorsal pericardial wall retain flexible fate potential and complex interactions with its neighboring cells during outflow tract formation^52,166^. Future efforts are warranted to resolve the detailed topology and cell-cell interactions during dorsal pericardial wall formation at the interface with the outflow tract.

The pericardium forms around the early-forming heart and accommodates an organ in constant motion. Significant research has focused on the inner-most pericardial layer, the epicardium, which envelopes the myocardium following pericardium formation and contributes to the heart in development as well as during homeostasis and regenerative responses^9,10,12,48,163,167–169^. As such, the epicardium exemplifies the tremendous tissue plasticity, organ-supportive properties, and disease involvement of mesothelial layers^9,10,12,20,22,23,48,164,170–172^. The dynamic, genetically undefined, and thin epithelial structure of the emerging pericardium among the complex ALPM presents challenges for a comprehensive investigation into its earliest development and subsequent specialization into pericardial and epicardial layers. Pericardial cells also pose challenges for live imaging even in optically accessible zebrafish embryos. Our light sheet-based tracking documented how pericardial progenitors migrate laterally and anteriorly over the yolk to encase the heart (**Fig. 2**), outside the field of view traditionally used to observe vertebrate heart formation *in vivo*^34,69,153,154,157,173,174^. Pericardial progenitors travel greater distances at higher speeds and with only few cell divisions compared to heart tube-forming progenitors (**Fig. 2**). Nonetheless, classic genetic insults to heart development including *mef2ca/b* perturbation^71,123^ and removing the entire endoderm as necessary for cardiac midline migration^126,128,130^ still permit pericardial progenitors to form its characteristic sac around even heavily deformed hearts (**Fig. 5**). These observations are in line with our model that the pericardium is indeed a mesothelial, and not a core cardiac lineage. In addition, our single cell transcriptome data indicates that mesothelial progenitors separate into distinct positional units in addition to the anterior-most pericardium (**Fig. 3B,E, Fig. 4A**). How pericardial progenitor migration and trajectories compare to other visceral mesothelial units forming around inner organs from the remaining progenitor cells warrants future investigation.

Our observed cell behaviors further underline the unique properties of pericardial cells in development that also carry over to post-natal stages at the transcriptional level (**Fig. 3G**). Due to unknown molecular definitions for their lineage and origins, and use of at times ambiguous naming as undefined mesenchyme, mesothelia in general remain underrepresented in the human or mouse transcriptome atlas^175,176^. Our single cell-based transcriptomics at the end of zebrafish gastrulation builds on our previous work that distinguished mesothelial progenitors among the emerging LPM as *hand2*-expressing populations^19^, expanded here into a more defined pericardial progenitor signature (**Fig. 3,4,6, SFig. 4,5**). Including *JAM2, SFRP5, TMEM88B, NR2F1, MEIS2*, and *TWIST1* as conserved core genes, this pericardial signature enables detection of pericardial and other mesothelial progenitors in publicly accessible, large single cell transcriptome datasets such as Zebrahub^76^ towards elucidating their lineage contributions and properties (**Fig. 4, SFig. 4, 5)**.

Integrating observations in zebrafish and neonatal rats, our data also expand our insights into the impact of canonical Wnt signaling on pericardium formation and tissue stiffness. Previous research in Xenopus discovered that Wnt/β-catenin signaling promotes proliferation in the pericardium while its inhibition is critical for differentiation of the myocardium^54,137,177^. In addition to the dynamic canonical Wnt/β-catenin activity in the ALPM and in pericardial/mesothelial progenitors (**Fig. 6**), we find that Wnt-inhibited embryos show a decrease in pericardial cells with resulting severe pericardial edema compared to controls, indicating that reduced Wnt signaling leads to fewer cells with increased individual cell size (**Fig. 7**). The canonical Wnt/b-catenin pathway has been repeatedly linked to cross-tissue interactions and features several secreted, negative feedback-associated signaling antagonists with powerful roles during development and homeostasis^54,133–135,178–183^. Relevant to pericardium formation, secreted Wnt antagonists of the SFRP family influence cardiogenesis, homeostasis, and disease^54,180,181,184–188^. In the early pericardial/mesothelial progenitors in zebrafish, we find *sfrp5* as prominently expressed gene among a core pericardial expression signature^19^ (**Fig. 3B,C,E**). In Xenopus, the paralog *sfrp1* expressed in cells lateral to the heart field has been shown to promote myocardial differentiation by inhibiting several Wnt ligand-driven responses^54^. Loss of *Sfrp1* in mice has been linked to progressive deterioration of heart function with increased fibrosis^187^. Notably, a cohort of pediatric patients with DCM presents with significantly elevated SFRP1 protein in their serum, which is sufficient to alter cardiomyocyte properties *in vitro*^150^. These observations indicate that persistent, pathogenic antagonism of canonical Wnt/β-catenin -catenin signaling might influence the heart but also the pericardium, with potentially detrimental impact on pericardial properties such as tissue stiffness or elasticity. In neonatal rats injected with recombinant sFRP1 and Iso to mimic the conditions of pediatric DCM^150,152^, we consistently found that isolated pericardia had significantly increased stiffness compared to controls (**Fig. 7K-L**). Together, these findings from zebrafish and rat models reveal that modulation of canonical Wnt/β-catenin signaling, through secreted antagonists like SFRP1, significantly affects pericardial cell composition and tissue rigidity. These data enhance our understanding of the developmental and pathological roles of Wnt signaling in pericardium formation and provides insights into the mechanical changes observed in pediatric dilated cardiomyopathy.

DCM is the most common form of cardiomyopathy and reason for cardiac transplantation in children and adults^189–193^. In pediatric DCM, five-year freedom from death or transplant remains low, ranging from 35%-40%^194^ in contrast, in adults with DCM five-year mortality ranges from 6.7% to 24.4%^195^. Effective treatments for adults have shown limited success in children^189–193^. Myocardial stiffness affects both diastolic and systolic function, which are common characteristics of pediatric heart failure^196^. While myocardial stiffness contributes to heart failure, pericardial stiffness as a potential factor in pediatric heart failure has not been thoroughly explored^197^. Our findings of increased pericardial stiffness upon persistent Wnt inhibition in our *in vivo* models (**Fig. 7**) proposes a possible contributing factor to pediatric DCM in patients with chronic, persistent exposure to SFRP1 as developmental Wnt antagonist^150^: the potential loss of pericardial elasticity counters efficient ventricle relaxation, aggravating the impaired function of the chamber. These insights have potential implications for the prognostic diagnosis and treatment options of pediatric DCM patients with elevated serum levels of Wnt antagonists. Taken together, conceptually incorporating the pericardium as mesothelial lineage into cardiogenesis provides a paradigm to connect its developmental and mechanical contributions to heart development, function, and disease.

## ACKNOWLEDGEMENTS

We thank C. Archer, A. Gilbard, and O. Gomez for zebrafish husbandry at CU Anschutz, Dr. Victor Ruthig for microscopy support, Greg Crouch and Robert Murchison for administrative support, and past and present members of the Mosimann lab for constructive experimental and conceptual input. We thank Drs. Caroline and Geoff Burns for kindly sharing *nkx2*.*5* transgenics, Dr. Stephanie Woo for *myl7* transgenics, Dr. Kristin Artinger for 7x*TCF:mCherry* reporter transgenics, and Dr. C. Ben Lovely for *sox32*/*cas* mutants. We thank Dr. Merlin Lange for discussions on using the Zebrahub data and Dr. Aaron Zorn for input on mesothelial anatomy and origins.

## METHODS

### Zebrafish Husbandry and Procedures

Zebrafish-related animal care and procedures were carried out in accordance with the veterinary office of the IACUC of the University of Colorado School of Medicine (protocol #00979), Aurora, CO, USA.

### Transgenic Zebrafish Lines and Transgene Activity

Established transgenic zebrafish lines used in this study include *TgBAC(hand2:EGFP)*^*pd24 61*^, *Tg(drl:mCherry)*^*zh705 62*^, *TgBAC(-36nkx2.5:ZsYellow)*^*fb7 33*^, *Tg(myl7:DsRed)*^*s879* 63^, and *Tg(7xTCF-Xla.Sia:NLS-mCherry)*^ia5 64^. Established mutant zebrafish lines used in this study include *hand2* (*han*^*S6*^)^65^ and *sox32* (*cas, sox32*^*ta56*^)^66^. The construct used to generate *Tg(tbx1:mCerulean)* was assembled from *pAF006* together with Tol2kit *#302* (*p3E_SV40polyA*), Tol2kit *pSN001* (*pMEminprom_mCerulean*) and *#394* (*pDestTol2A2*) as backbone^67–69^. Gateway cloning reactions were performed with the Multisite Gateway system with LR Clonase II Plus (Cat#12538120; Life Technologies) according to the manufacturer’s instructions and concentration calculations^70^. Founder lines were screened for single integrations and compared to the existing *Tg(tbx1:EGFP)*^zh702^ zebrafish line^69^ to ensure faithful expression.

### Morpholino Injections

The previously characterized and validated *mef2ca* and *mef2cb* morpholinos^71^ (*MO1-mef2ca*^*ATG*^:*5’-TTTCCTTCCTCTTCCAAAAGTACAG-3’*, 1.0 ng; *MO2-mef2cb*^*ATG*^:*5’TGTCCCCGTCTTTTCGTCTCTCTCT-3’*, 0.25 ng) were obtained by GeneTools, LLC and injected into the one-cell stage of *hand2:EGFP* transgenic embryos. The MOs were kept in stock of 1 mM and diluted prior to injection into the yolk of 1-cell stage zebrafish embryos at ∼2 nl.

### Single cell RNA-Sequencing

Around 1000 *drl:mCherry;hand2:EGFP* double-transgenic embryos were grown at 28.5 °C in E3 medium until tailbud stage (10 hpf) was reached. Embryos were dechorionated by incubating in 1 mg/mL Pronase (Sigma, 53702-50KU) and rinsed with E3 medium. For dissociation, PBS was replaced with 2 mg/mL collagenase IV (Worthington) in DMEM (high glucose (4.5 g/l) and NaHCO3, without L-glutamine and sodium pyruvate, Sigma-Aldrich). Embryos were incubated for 2x 5 min in a water bath at 37°C and carefully pipetted up and down into a single-cell suspension. Cells were filtered through a 35 μm cell-strainer (Falcon, round-bottom tubes with cell-strainer cap) and centrifuged at 400xg for 30 sec. Cells were washed in 1X HBSS (Gibco) containing 2% FBS. After washing, the cells were centrifuged again, and the pellet was resuspended in 1X PBS. Next, cells were sorted to isolate EGFP- and mCherry-expressing cells using the MoFlo XDP100 sorter at the CU-SOM Cancer Center Flow Cytometry shared resources platform at CU Anschutz Medical Campus. Sorted cells were collected in a 1.5 mL FBS-coated microcentrifuge tube.

### Bulk RNA-sequencing

RNA was extracted from tissues using the mirVana kit (Ambion) and reverse transcribed into complementary DNA using the iScript cDNA Synthesis Kit (Bio-Rad)^72^. Samples with RNA integrity above 9 RIN were considered suitable and high quality for RNA-seq. Bulk RNA-seq was performed by the University of Colorado Genomics Core as extensively described by our group. 1X150 directional mRNA sequencing was performed using an Illumina NovaSeqX resulting in an average of 40 million mapped reads per sample. Bulk RNA-seq sequencing were mapped to Rnor_6.0 and genes quantified using the nfcore rnaseq v3.12.0 pipeline (https://zenodo.org/records/10171269). Fragments per kilobase of exon per million mapped reads values were calculated using Cufflinks for each sample (*n* = 5 pericardial tissue, *n*=7 myocardial tissue (6 males and 1 females). Differentially expressed genes were calculated using the nfcore differentialabundance v1.3.1 pipeline (https://zenodo.org/records/10046399) starting with the raw counts file.

### Metascape Analysis

Using Metascape^73^, we first identified all statistically enriched GO/KEGG terms for the top 50 genes in each respective cluster, accumulative hypergeometric p-values and enrichment factors were calculated and used for filtering. The remaining significant terms were then hierarchically clustered into a tree based on Kappa-statistical similarities among their gene memberships. Then, a 0.3 kappa score was applied as the threshold to cast the tree into term clusters.

### Single-Cell Analysis

10X Genomics Chromium technology was used to capture and profile single cell transcriptome 3’ gene expression (Genomics Core at CU Anschutz). Generated libraries were sequenced on the Illumina NovaSeq 6000 instrument at the University of Colorado Cancer center. Upon sequencing, Fastq sequencing files from were processed through Cell Ranger (v5.0.1)^74^ with a zebrafish *GRCz11* library to obtain UMI gene expression counts. These were analyzed using the standard methods in the Seurat pipeline^75^. After clustering at a resolution of 0.8 snn, the mesothelium cluster was sub-clustered to separate the mesothelium and pericardium clusters. Clusters were annotated manually using marker gene expression and the differentially expressed genes for each cluster were calculated with the FindAllMarkers function. The Zebrahub zebrafish embryonic single cell RNA-seq data was downloaded from https://zebrahub.ds.czbiohub.org^76^. Annotated primitive heart tube/heart cells from 12 hpf, 14 hpf, 16 hpf, and 19 hpf scRNA-seq were subset and re-analyzed to generate a UMAP containing pericardium, myocardium and precursor clusters. The PCA reduction of this subset was used to generate pseudotime lineages using Slingshot^77^.

### Gene Expression Analysis and *in situ* Hybridization (ISH)

First-strand complementary DNA (cDNA) was generated from zebrafish RNA isolated from 18 hpf zebrafish embryos using SuperScript III first-strand synthesis kit (Invitrogen). DNA templates were generated using first-strand cDNA as a PCR template and the primers as specified for each gene of interest; for in vitro transcription initiation, the T7 promoter *5′-TAATACGACTCACTATAGGG-3′* was added to the 5′ ends of reverse primers. Specific primers used were *jam2b*: forward: *5’-CTAACCTCTGCTCTTCTC-3’* reverse: *5’-TAATACGACTCACTATAGGGCATTGTCATGTTCAGCTC-3’* and *sfrp5*: forward: *5’-GAATCACAGCAGAGGATG-3’* reverse: *5’-TAATACGACTCACTATAGGGCATCTGTACTAATGGTCG -3’*. PCR reactions were performed under standard conditions as per manufacturer’s protocol using Phusion high-fidelity DNA polymerase (Thermo Fisher Scientific). RNA probes were generated via overnight incubation at 37 °C using 20 U/µL T7 RNA polymerase (Roche) and digoxigenin (DIG)-labeled dNTPs (Roche) as per manufacturer’s protocol. The resulting RNA was precipitated in lithium chloride and EtOH. Wildtype strain zebrafish embryos were fixed in 4% PFA in PBS overnight at 4 °C, dechorionated, transferred into 100% MeOH, and stored at −20 °C. ISH of whole-mount zebrafish embryos was performed essentially as per standard protocol in the field^19,78^ and compared to publicly available data on ZFIN^79^ when possible.

### Microscopy and Image Analysis Confocal Imaging

Embryos were anesthetized at 3 dpf with 0.016% Tricaine-S (MS-222, Pentair Aquatic Ecosystems, Apopka, FL, USA, NC0342409) in E3 embryo medium. Laser scanning confocal microscopy was performed on a Zeiss LSM880 following embedding in E3 with 1% low-melting-point agarose (LMA) (Sigma-Aldrich, A9045) on glass bottom culture dishes (Greiner Bio-One, 627861). Heartbeat was stopped with 50 mM 2,3-butanedione monoxime (BDM, Cat#B0753; Sigma) as indicated in individual experiments. Images were collected with a x10/0.8 air-objective lens, Plan-Apochromat ×20/0.8 M27, or with Plan-Apochromat 40x/1.3 Oil DIC M27 objective lenses. All channels were captured sequentially with maximum speed in bidirectional mode, with the range of detection adjusted to avoid overlap between channels. Maximum projections of acquired Z-stacks were made using ImageJ/Fiji^80^ (2.14.0), cropped and rotated using Adobe Photoshop 2024 (24.7.0), and assembled in Adobe Illustrator (27.8.1).

### Light sheet Imaging

Embryos used for long-term imaging were treated with 0.003% 1-phenyl-2-thiourea (PTU, Sigma-Aldrich) to prevent melanin pigment formation. The Zeiss Z.1 microscope equipped with a Zeiss W Plan-Apochromat 20×/0.5 NA objective was used for all other light sheet microscopy^19,69,81,82^ and as mentioned in the figure legends. Embryos were embedded out of the chorion in 1% LMA respectively a 50 or 20 µL glass capillary. Live embryos older than 16 ss were additionally mounted with 0.016% ethyl 3-aminobenzoate methanesulfonate salt (Tricaine, Sigma-Aldrich) in the LMA and added to the E3 medium to prevent movement during imaging. For the multi-angle imaging data sets: we manually registered the four individual angles per embryo and then applied the Fiji *Multiview Reconstruction* and *BigStitcher* plugins for fusion and deconvolution of the images^83–86^. Images and movies were further processed using ImageJ/Fiji^80^ (2.14.0) and Imaris (9.7.2). The Mercator projection in Figure 1E is derived from our previous live imaging datasets ^19,81^.

### Cell Tracking Analysis

Cell tracking was performed using Imaris’ Object Classification and Lineage tools, as extension of previously reported manual backtracking of developmental cell lineages^87^. The 3D image data were first processed using Imaris’ segmentation tools to identify the developing heart field as the region of interest. The Object Classification tool was used to define cells as either ‘pericardial’ or ‘cardiac’ based on their size, shape, and intensity characteristics. The Lineage tool was employed to establish and track the lineage of cells over time by linking cells across different time points to monitor their movement, division, and fate. Tracking parameters such as search radius and maximum displacement were adjusted to optimize accuracy, with manual corrections applied to address any tracking errors or inconsistencies. Lineage trees were generated, and individual cell trajectories were reviewed to analyze cell division patterns and migration paths. Data analysis was conducted using Imaris’ quantitative tools to measure cell proliferation, migration, and spatial relationships. Metrics such as cell velocity and division rate were extracted and compared. Various visualization options, including 3D and 4D renderings, were employed to illustrate key findings. Validation of the tracking results was performed by cross-verifying with manual observations or alternative methods to ensure accuracy and reliability.

### Chemical Treatments

IWR-1-endo (Sigma-Aldrich, 681669) was administered to embryos at 10 µM in E3 at the 18 ss (18 hpf equivalent) overnight and washed out through several washing steps of E3. 2,3-Butanedione monoxime (2,3 BDM, Thermo-Fisher, A14339-22) was administered to embryos at 10 mM in E3 at the 18ss (18 hpf) overnight and washed out through several washing steps of E3. Embryos were then raised in E3 to 3 dpf to visualize the pericardial cavity. Controls were treated with an equivalent amount of DMSO. Cell density was quantified using maximum intensity projections in Fiji software by drawing a region of interest (ROI) around the pericardium (labeled by *hand2:EGFP*) and measuring the number of GFP+ nuclei within the area. Images were blinded prior to quantification, and cell size was quantified by measuring the length of the widest point of 6 cells across at least 9 embryos per condition (3 independent experiments).

### Antibody Staining

Embryos were fixed in 4% formaldehyde, 0.1% TritonX in PEM (0.1 M PIPES, 2 mM MgSO4, and 1 mM EDTA)^69^ for 2 hours at room temperature, washed in 0.1% PBS TritonX (PBSTx), and permeabilized in 0.5% PBSTx. Blocking was done in blocking buffer containing 5% goat serum, 5% BSA, 20 mM MgCl2 in PBS, and embryos/hearts incubated with primary antibodies diluted in blocking buffer at 4 °C overnight. anti-MHC primary antibody (MF20, 53-6503-82, Invitrogen, 1:50) was used with the Alexa-conjugated 594 secondary antibody (A-11012, ThermoFisher, 1:500) in 0.1% PBSTx at 4 °C overnight. Before imaging, embryos were mounted in 1% low-melting-point agarose.

### Statistics

Unpaired non-parametric (Mann-Whitney) two-tailed t-test was done to compare the scores between two groups. For analyses with more than two groups, 1-Way ANOVA was performed to compare the scores between the groups. Adjusted p-values after multiple tests correction are reported and significance was set at p < 0.05. Quantifications of cell size and density were performed blind.

### Neonatal rat injections

All animal studies were approved by the Institutional Animal Care and Use Committee (IACUC) of the University of Colorado Anschutz Medical Campus (protocol #00527), Aurora, CO, USA. Pregnant Sprague Dawley female rats were purchased from Charles Rivers laboratories. All animals were housed in the animal facility of the University of Colorado Anschutz Medical Campus and monitored daily. Male and female young (0-5 day old) Sprague Dawley rats were treated with 100µg/kg/day Isoproterenol, 50µg/kg/day sFRP1 and vehicle treated controls (PBS, 0.5mM ascorbic acid) every day for 5 days and every other day for 7 days by intraperitoneal injection. The sFRP1 (Recombinant Human sFRP-1 Protein, CF, R&D systems) solution was freshly prepared for each treatment by dissolving in phosphate-buffered saline (PBS) at room temperature. Isoproterenol (Isoproterenol hydrochloride, Sigma Aldrich St. Louis, MO) was freshly prepared and dissolved in 0.5mM of ascorbic acid water.

### Isolation of myocardium and pericardium

At the end of the study period, rats were euthanized. Myocardial and pericardial tissue were isolated, immediately weighed, frozen in liquid nitrogen and stored at -80 °C for further tissue analysis.

### AFM assessment for pericardial stiffness

Frozen pericardial tissue were embedded in optimal cutting temperature (OCT) (Sakura) and cryosectioned at 5 microns one section per slide. Tissues were allowed to equilibrate at cryostat temperature of -20°C prior to cryosectioning. Isolated pericardial tissue on slides were monitored, selected and their morphological details observed with an optical light microscope. The physical and physiological cues regarding sample preparation and AFM analysis were kept constant across all tissue samples. Three (3) pericardial tissue per treatment group were analyzed. The methodology for AFM measurement and analysis was based on our previous studies^88–91^. Briefly, pericardial stiffness was determined using a NanoWizard® 4a (JPK Instruments, Carpinteria, CA, USA). In quantitative imaging (QI) mode with a qp-BioAC-1 (NanoandMore, Watsonville, CA, USA) cantilever. The cantilever force constant was in the range of 0.15 to 0.55 N/m. Calibration of the cantilever was performed using the thermal oscillation method prior to each experiment. A 5,625 µm^2^ area was scanned using a set point of 5 nN and a Z-length of 2 µm. Several pericardial orientations were scanned across the sample, and they were considered for the mechanical average of the sample. Four random scans were performed per sample. Every scan was composed with over 60,000 force curves (60,000 nanomechanical data points per scan). The Hertz model was used to determine the mechanical properties of the pericardium using the JPK software and a correction for an offset in the height data was performed line by line using the JPK data processing operation.

## Data availability

All scRNA-seq analyses were run through Cell Ranger (v5.0.1)^74^ with a zebrafish *GRCz11* library and analyzed using the Seurat 5 R pipeline^75^, and is browsable at https://cuanschutz-devbio.shinyapps.io/Moran_scRNAseq/. Bulk RNA-seq sequencing were mapped to Rnor_6.0 and genes quantified using the nfcore rnaseq v3.12.0 pipeline (https://zenodo.org/records/10171269). Differentially expressed genes were calculated using the nfcore differentialabundance v1.3.1 pipeline (https://zenodo.org/records/10046399) starting with the raw counts file, and are browsable at https://cuanschutz-devbio.shinyapps.io/Moran_rat_bulkRNAseq/. The Zebrahub zebrafish embryonic single cell RNA-seq data^76^ was downloaded from https://zebrahub.ds.czbiohub.org. The PCA reduction of this subset was used to generate pseudotime lineages using Slingshot^77^. Code for all sequencing data included in this manuscript are available at https://github.com/rebeccaorourke-cu/Moran_scRNAseq_manuscript. Code of the custom-made processing steps for light sheet imaging^86^ are available at https://github.com/DaetwylerStephan/multi_sample_SPI <u>M</u>.

---

**Supplementary Figure 1:**
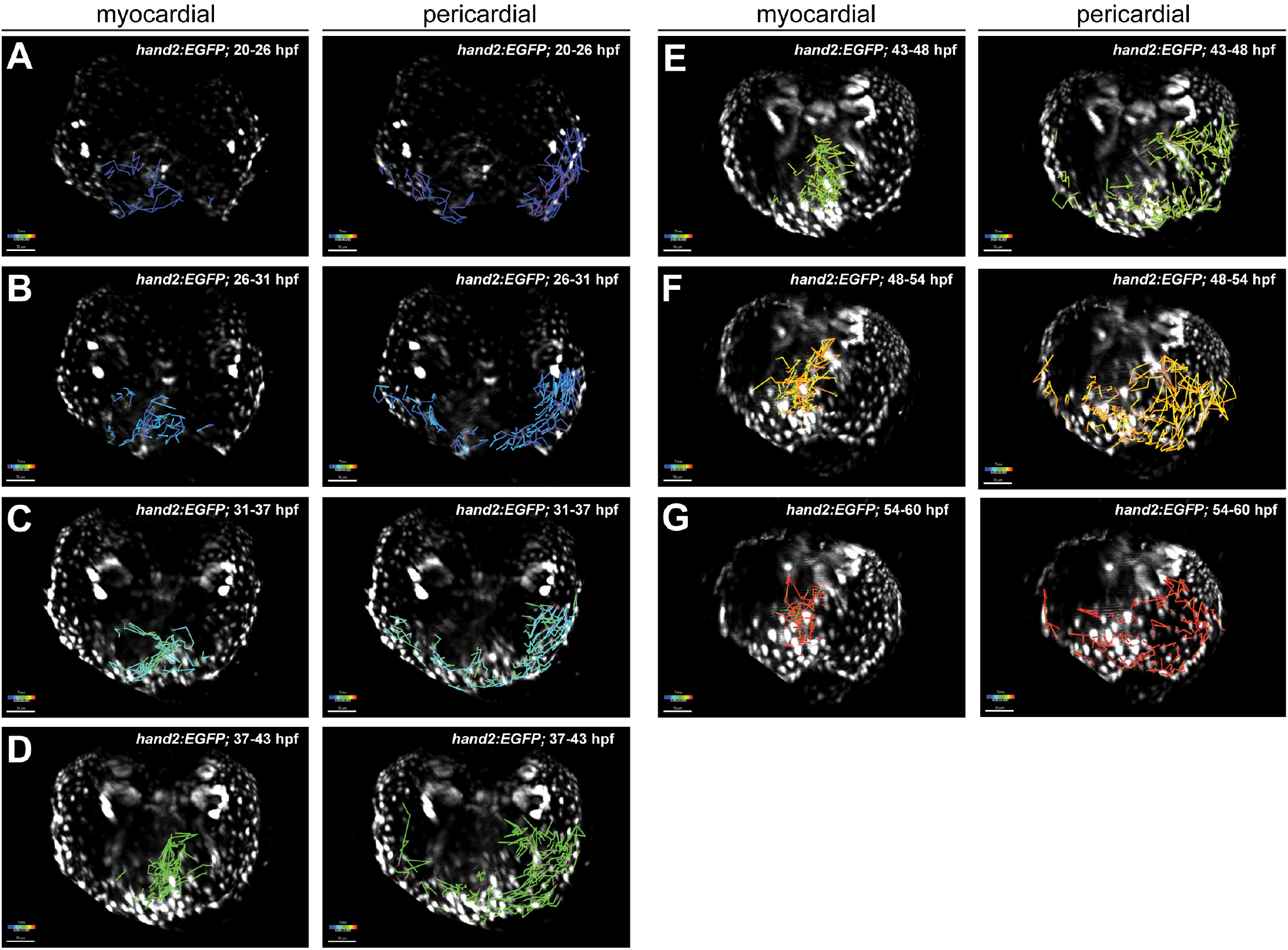
Individual lineage tracks of myocardial and pericardial cells. **A**.Tracks of *hand2:EGFP* myocardial (left) and pericardial (right) lineage cells from 20 to 26 hours post-fertilization (hpf), **B**.26 to 31 hpf, **C**. 31 to 37 hpf, **D**. 37 to 43 hpf, **E**. 43 to 48 hpf, **F**. 48 to 54 hpf, and **G**. 54 to 60 hpf. Anterior to the bottom, see also Figure 2.

**Supplementary Figure 2:**
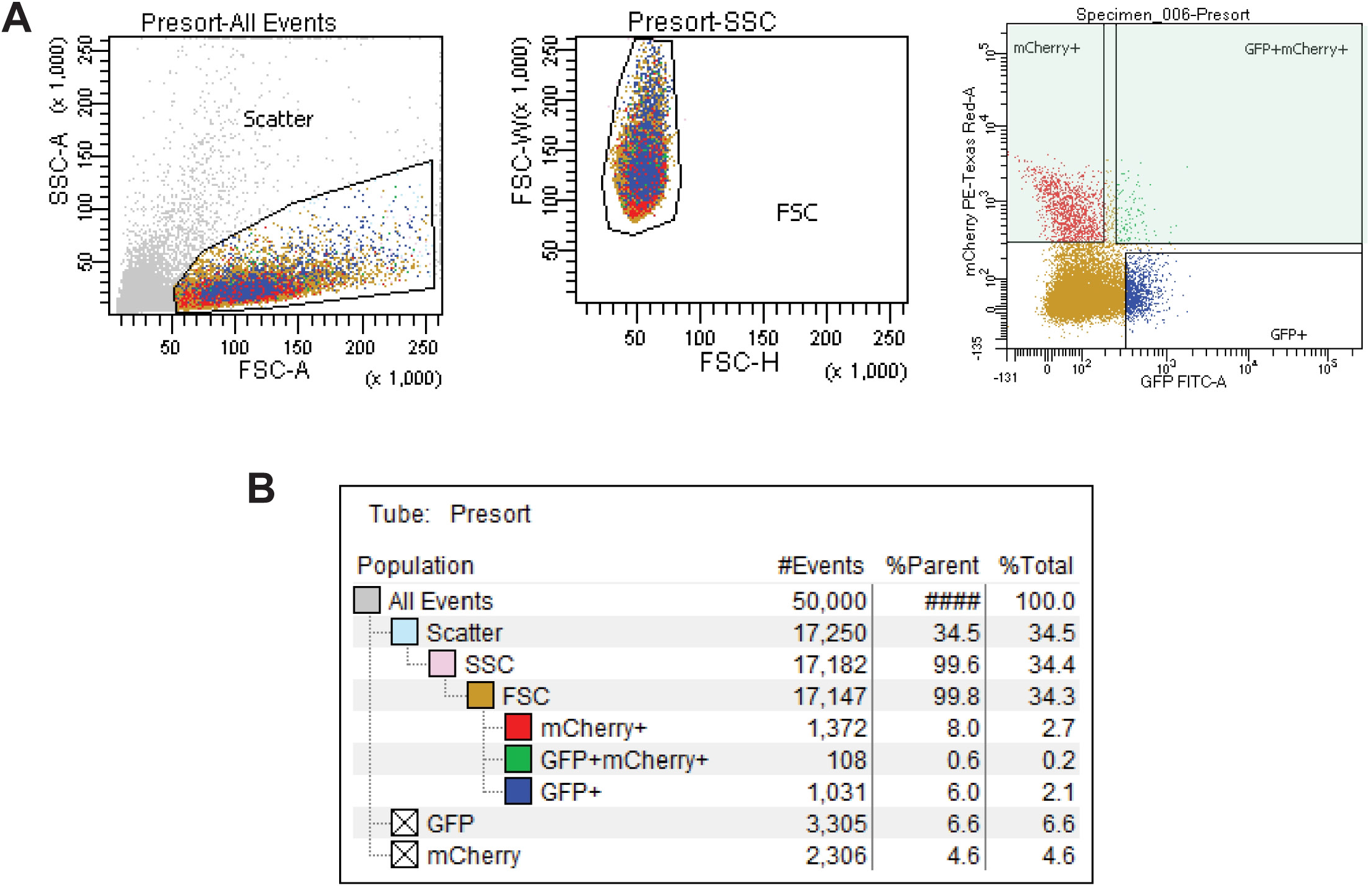
FACS for mCherry signal from *hand2:EGFP;drl:mCherry* embryos at 10 hpf. **A**. FACS plots with gating and sorting strategy for mCherry-positive cells within the single-cell suspension of *hand2:EGFP;drl:mCherry*-dissociated embryos at tail bud stage (approx. 10 hpf). **B**. Table with the percentage of mCherry- and EGFP-positive cells in the total amount of cells. See also Figure 3.

**Supplementary Figure 3:**
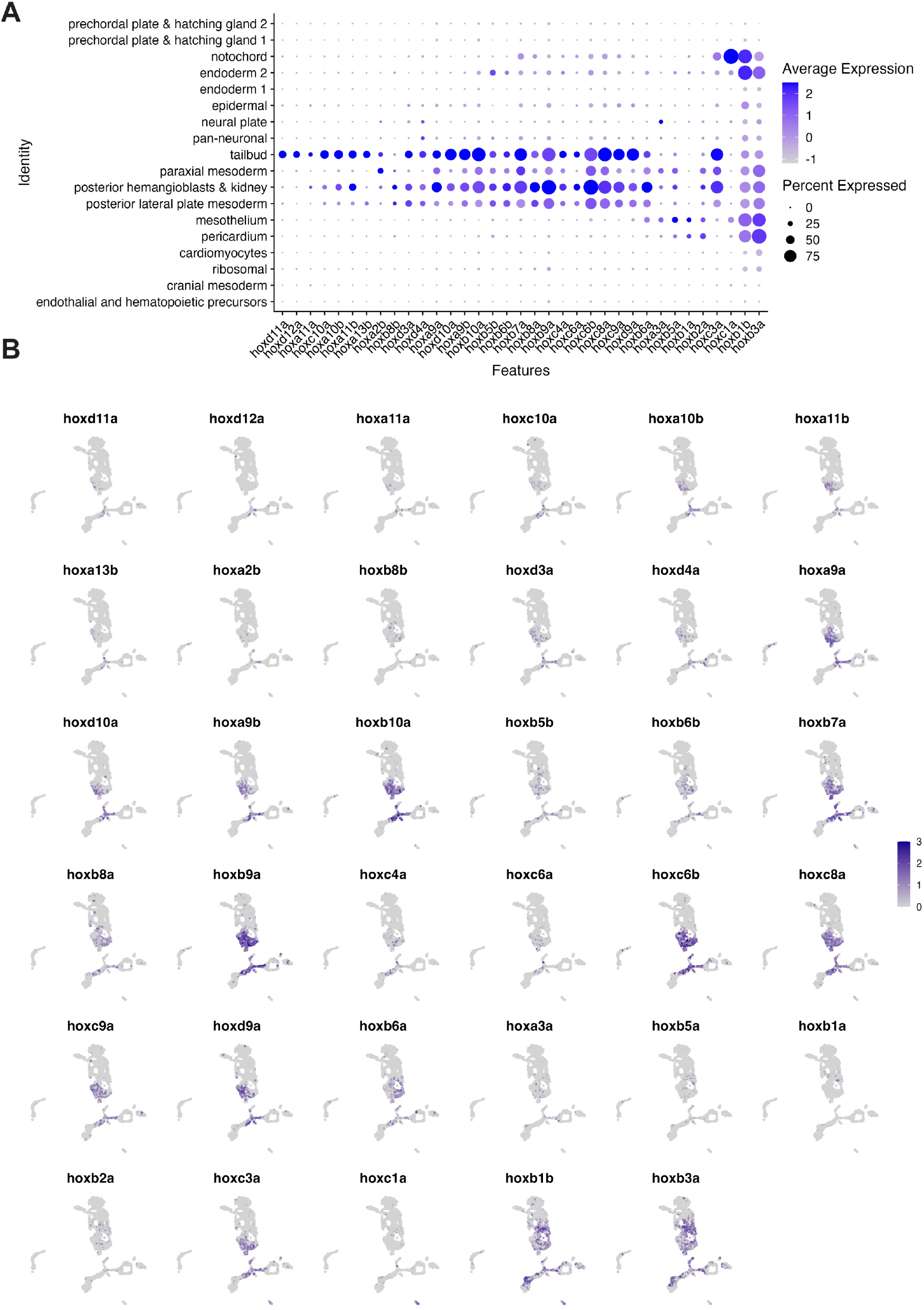
Distribution of *hox* gene expression across LPM clusters. **A**. Dotplot including all *hox* genes expressed throughout our designated 18 clusters. Dots are colored by column-scaled mean expression (log-transformed library-size-normalized counts) and sized by detection frequency (fraction of cells with non-zero counts); rows and clusters are ordered according to hierarchical clustering of scaled expression values. **B**. UMAP plots of all the *hox* genes expressed throughout the data set. Cells are colored by scaled expression values using top and bottom 1%-quantiles as boundaries.

**Supplementary Figure 4:**
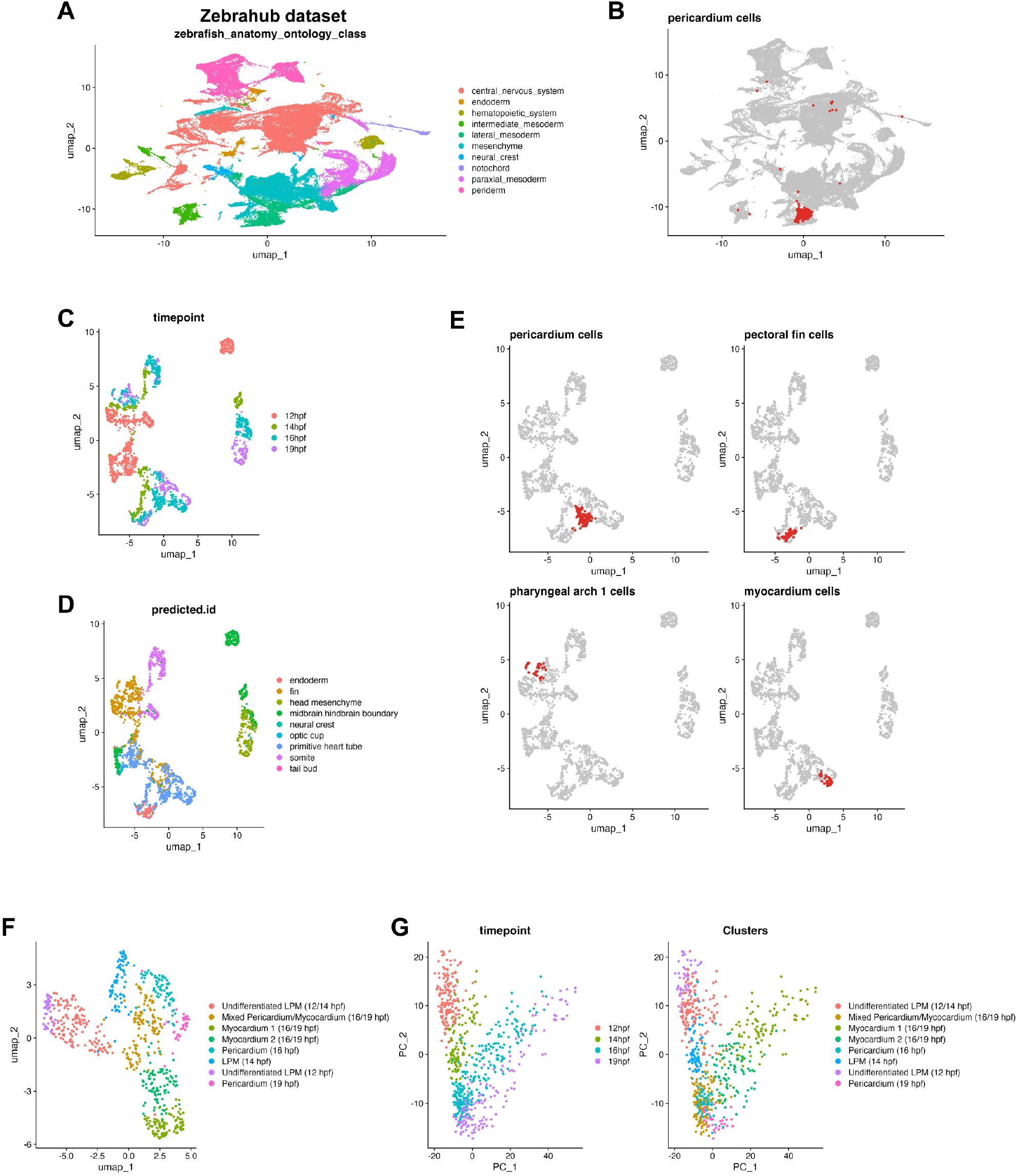
Zebrahub scRNA-seq dataset to subset pericardium and myocardium cells. **A**. UMAP projection of zebrafish cells in Zebrahub dataset, colored by anatomical ontology classes, highlighting the diversity of cell types within the embryo. **B**. UMAP projection of zebrafish embryonic cells with pericardium cells highlighted in red, showing their distribution across different clusters. **C**. UMAP projections highlighting specific cell types within the zebrafish embryo: pericardium cells, pectoral fin cells, floor plate cells, pharyngeal arch 1 cells, pharyngeal arch 2 cells, and myocardium cells, each marked in red to show their spatial organization within the clusters. **D**. UMAP projections of zebrafish embryonic cells, with cells colored by cluster (left) and by time point (right), demonstrating the developmental stages and the organization of clusters over time. **E**. UMAP projection of a subset of clusters identified in zebrafish embryonic cells, highlighting the specific lineage groups: undifferentiated lateral plate mesoderm (LPM) at different hours post-fertilization (hpf), pericardium, and myocardium cells. **F**. Principal component analysis (PCA) of zebrafish embryonic cells, showing the distribution of cells over time (time point). **G**. PCA projection of the subset clusters identified in **F**, demonstrating the developmental trajectory of these specific lineages over time. See also Figure 4.

**Supplementary Figure 5:**
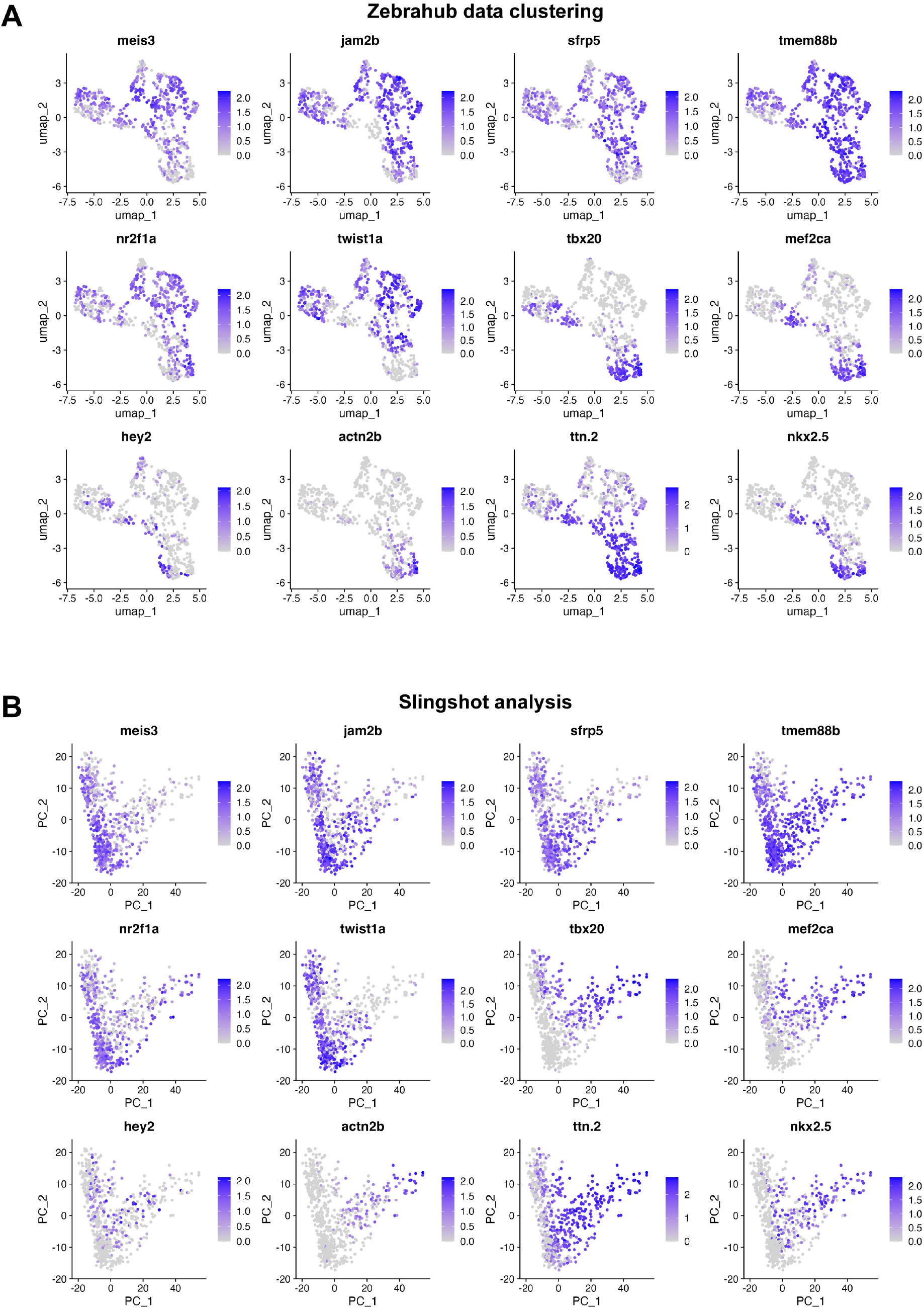
Gene expression of pericardial and myocardial genes from Zebrahub sub-setting. **A**. UMAP projections of representative genes found among LPM, myocardial, and pericardial clusters from zebrafish embryonic cells across time as key cell fate markers used to annotate clusters. Cells are colored by scaled expression values using top and bottom 2%-quantiles as boundaries. **B**. PCA projections of LPM, myocardial, and pericardial clusters from zebrafish embryonic cells across time showing key cell fate markers used to annotate clusters. Cells are colored by scaled expression values using top and bottom 2%-quantiles as boundaries. See also Figure 4.

